# Reduced Model-Based Control in Gambling Disorder Despite Intact Neural Value and Task Structure Representations

**DOI:** 10.1101/2025.07.18.665478

**Authors:** A. Brands, K. Knauth, D. Mathar, S. Lee, B. Kuzmanovic, M. Tittgemeyer, J. Peters

## Abstract

Disordered gambling has been linked to impairments in goal-directed (model-based) control and reinforcement learning. Here we investigated the potential neural basis of this impairment using a sequential reinforcement learning task (modified two-step-task), computational modeling, and functional magnetic resonance imaging (fMRI) in individuals exhibiting symptoms of disordered gambling (GD) and matched healthy controls (HC, n=30 per group). Model-agnostic analyses replicated the effects of reduced performance and reduced model-based control in the gambling group, both in terms of choice and response time effects. Computational modeling of choice behavior confirmed that this effect was due to reduced model-based control in the gambling group. Analyses of choices and response times using drift diffusion modeling revealed a more complex pattern, where behavioral impairments in the gambling group were linked to changes across several parameters reflecting drift rate modulation and asymptote, as well as non-decision time. Despite these pronounced behavioral differences, the gambling group exhibited largely intact neural effects related to the task transition structure, reward feedback and trial-to-trial behavioral adjustments. Results are discussed with respect to current neurocomputational models of behavioral dysregulation in disordered gambling.

## Introduction

Gambling disorder (GD) is a behavioral addiction that can have detrimental effects on quality-of-life indices, including personal finances, work, relationships and overall mental health (Blaszczynski & Nower, 2002; Muggleton et al., 2021). Accumulating evidence suggests similarities of gambling disorder and substance-use-disorders (SUDs) both on behavioral, cognitive and neural levels (Balodis & Potenza, 2020; Fauth-Bühler, et al., 2016; Leeman & Potenza, 2012; Lobo et al., 2015; Petry, 2010; Singer et al., 2020). In light of these similarities, the fifth edition of the “Diagnostic and Statistical Manual of Mental Disorders” categorizes gambling disorder in the category of “Substance-related and Addictive Disorders” (American Psychiatric Association, 2013). In contrast to SUDs, differences in behavioral and/or neural effects between individuals with GD and controls are unlikely to be confounded by chronic or acute drug effects (Clark et al., 2019; Fauth-Bühler, et al., 2016; Leeman & Potenza, 2012). Gambling disorder has thus been termed a “pure addiction” (Dixon et al., 2006).

Categorical definitions of mental disorders have increasingly been called into question. The National Institute for Mental Health of the United States proposed the Research Domain Criteria (RDoC) to foster characterization of the dimensions underlying mental disorders. According to RDoC, research should focus on the identification of continuous neuro-cognitive dimensions that might go awry in mental disorders, i.e. trans-diagnostic markers (Nelson et al., 2016). One promising candidate for such a trans-diagnostic process that is affected across a range of psychiatric conditions is model-based control during sequential reinforcement learning (Daw et al., 2011; Voon et al., 2017). Model-based control refers to computationally more expensive goal-directed strategies that utilize models of the environment, contrasting with model-free (MF) behavioral control that operates on stimulus-response associations (Balleine & O’Doherty, 2010; Daw et al., 2011; Doll et al., 2012; Voon et al., 2017). Changes in the contributions of model-free and model-based control have been reported in schizophrenia (Culbreth et al., 2016), obsessive compulsive disorder (Gillan et al., 2020), SUDs (Sebold et al., 2014; Ruan et al., 2024; but see Magrabi et al., 2022) and GD (Wyckmans et al., 2019; Bruder et al., 2021). Reduced model-based control is also associated with sub-clinical psychiatric symptom severity (Gillan et al., 2016).

On the neural level, model-free and model-based control have been linked to activation in partly overlapping yet distinct key structures within the mesolimbic/mesocortical dopaminergic system, including the ventral and dorsal striatum, as well as ventromedial and lateral prefrontal cortex (Daw et al., 2011; Doll et al., 2015; Lee et al., 2014; Huang et al., 2020). Functional changes in these circuits have been linked to SUDs (Reiter et al., 2016; Chen et al., 2021). Yet, it is unclear if similar functional changes relate to reductions in model-based control in individuals suffering from GD.

Our first aim was therefore a replication of attenuated model-based control during sequential reinforcement learning in GD (e.g., Wyckmans et al., 2019; Bruder et al., 2021). We next leveraged hierarchical Bayesian modeling using reinforcement learning drift diffusion models (RLDDMs) (Fontanesi et al., 2019; Peters & D’Esposito, 2020; Wagner et al., 2022) to elucidate computational mechanisms underlying the predicted performance deficit in GD. Functional magnetic resonance imaging (fMRI) was used to probe circuits previously implicated in reward valuation and model-based control for potential group differences during task performance.

## Methods

### Participants

At the time of study planning, no data on model-based control in GD were published. We therefore based our power analysis based on the well-replicated effect of steeper reward discounting in GD compared to controls (Miedl et al. (2012), Cohen’s *d* = 0.67). With power of 0.8 and α = 0.05, this led to a target sample size of n=30 per group. An all-male sample of n=30 individuals exhibiting problematic gambling behaviour (≥2 DSM-5 criteria, GD group) and n=30 healthy male controls (HC) participated. Exclusion criteria were current or prior psychiatric or neurological diseases, current medication or drug abuse, as well as not being fluent in German. Participants were recruited via local online bulletin boards.

Participants were pre-screened to confirm regular participation in gambling. Subsequently, a semi-structured clinical interview was conducted, and participants from the GD group met at least two DSM-5 criteria for GD. Groups were matched on age, education, and socioeconomic status. In addition to the DSM-5 criteria, gambling severity was assessed using the the “Kurzfragebogen zum Glücksspielverhalten” (KFG; Petry & Baulig, 1996), and a German adaptation of the South Oaks Gambling Screen (SOGS; Lesieur & Blume, 1987).

As planned, the final sample consisted of n=30 individuals who exhibiting at least two DSM-5 symptoms of disordered gambling (GD) and n=30 matched healthy controls (HC, mean age ± SD; GD = 28.63 ± 6.37; HC = 27.27 ± 6.15). On average, subjects in the GD group fulfilled 5.47 DSM-5 criteria (SD=2.32, range 2 to 9). All participants were right-handed and without a history of psychiatric or neurological illness nor current medication intake. A negative drug screening was confirmed via a urine drug test prior to participation in the main testing session.

All procedures were approved by the ethics committee of the University of Cologne Medical Center (application number: 17-045), and participants provided informed written consent prior to participation.

### Procedure

Data collection took place at the Imaging Center of the Max Planck Institute for Metabolism Research in Cologne, Germany. Upon arrival, participants first gave written informed consent, were assessed regarding height and weight, performed a urine drug screening, and completed a short questionnaire on their current condition. Participants were then given detailed instructions about the sequential reinforcement learning task they would perform in the fMRI environment. They were informed that their goal was to maximize their monetary earnings during the task. Before entering the MRI scanner, participants completed 20 training trials. This was followed by 300 trials conducted in three separate sessions inside the scanner. Following the MRI sessions, participants completed an implicit learning task (Weather Prediction Task; Knowlton, 1994) and three working memory tasks (operation span (Redick et al.,2012), listening span (van den Noort et al., 2008), and digit span (Wechsler, 2008). Finally, they filled out a computerized questionnaire battery, which included assessments of gambling disorder severity (DSM-V, Falkei & Wittchen, 2015) and gambling-related cognitive distortions (South Oaks Gambling Screen, SOGS, Lesieur & Blume, 1987; Kurzfragebogen zum Glücksspielverhalten, KFG, Petry & Baulig, 1996; and Gambling Related Cognitions Scale, (GRCS, Raylu & Oei, 2004), impulsivity (Barratt Impulsiveness Scale, BIS-15)(Spinella, 2007), depressive symptoms (Beck’s Depression Inventory, BDI-II)(Beck et al., 1996), as well as nicotine (Fragerström Test for Nicotine Dependence; FTND, Fragerström & Schneider, 1989), and alcohol dependence (Alcohol Use Disorders Identification Test; AUDIT, Saunders et al., 1993).

Additionally, we computed a compound score of gambling disorder severity (GD score) by averaging z-scores across DSM-V, SOGS, and KFG.

Clinical and demographic measures (BDI-II-, FTND-, and AUDIT-scores, age) were included in the second level fMRI analysis as covariates (see below). GRCS and addiction severity scores and clinical measures were additionally used in regression models to predict individual differences in model parameters (see below).

### Two-Step Reinforcement Learning Task

Participants performed a modified version of a sequential reinforcement learning task (two-step task; TST) (Daw et al. (2011)) during fMRI. The task was programmed using the Psychophysics Toolbox Version 3.52 implemented in MATLAB R2014b software (The Mathworks Inc., MA, USA). Following Kool et al. (2016), the outcome stage used drifting reward magnitudes (Gaussian random walks with boundaries at 0 and 100) instead of drifting reward probabilities, as in the classical task version (Otto et al., 2013; Daw et al. 2011). The number of trials was increased to 300 to improve model estimation. Trials were separated by inter-trial intervals sampled from a Poisson distribution (M = 2 sec, range: 1–9 sec). Two 30-second breaks were included after 100 and 200 trials.

Each trial consisted of two stages: in the first stage (S1), participants chose between two fractals embedded in grey boxes. After this first-stage choice (response time limit: 2sec), they proceeded to one of two second stages (green or blue), according to the task transition structure. S1options were probabilistically linked to transitions to one of the second stages (S2; 70% common transition vs. 30% rare transition, vice versa for the other S1option). InS2, participants again chose between two new fractals in coloured boxes. Following the second -stage choice (response time limit: 2sec), participants were presented with the trial outcome, which varied between 0 and 100 cents (2sec).

To optimize reward accumulation, participants had to learn two key aspects of the task. First, they had to learn the transition structure, i.e. specifically, which first-stage choice was more likely (70% of trials, common transition) to lead to which second -stage (green or blue). Second, they had to track the fluctuating reward pay-offs linked to each S2 option. Pay-offs followed a Gaussian random walk with gradual changes and reflecting boundaries at 0 and 1 (which were converted to a 0-100 range for display in the task). Participants received explicit instructions regarding the task structure and completed twenty practice trials with a separate set of random walks and stimuli (to avoid transfer effects) prior to participation in the fMRI experiment to familiarize participants with the procedure.

### Data Analysis

Behavioural data from the TST task were analysed using two complementary approaches, *model-agnostic* and *model-based*. Model-agnostic analyses used standard linear mixed models, whereas model-based analyses used hierarchical Bayesian reinforcement learning models.

#### Model-Agnostic Analyses

Model-agnostic analyses focused on two outcome measures: (1) the probability of repeating the same first-stage choice as on the previous trial (stay probability, *p*(stay)), and (2) response times (RTs) at both the first and second stage. The *p*(stay) analysis assesses participants’ reliance on model-free versus model-based learning by examining how first-stage choice repetitions depend on the previous trial’s reward outcome (high reward vs. low reward) and the transition type (common vs. rare). High vs. low reward trials were classified by comparing the outcome received on the current trial to the mean reward obtained over the last twenty trials. A main effect of reward, with increased *p*(stay) after high reward trials irrespective of transition is thought to reflect model-free behaviour driven by reinforcement (Daw et al., 2011; Otto et al., 2013; Sutton & Barto, 2018; Voon et al., 2017).In contrast, a reward × transition interaction is generally assumed to reflect model-based control, since participants utilize an internal model of the task transition structure (Daw et al., 2011; Otto et al., 2013).

We fitted linear mixed models (LMMs), testing for main effects of reward, transition, and group (GD vs. HC), as well as their interactions. To control for potential confounds, BDI, BIS, FTND, and AUDIT scores, along with any demographic variables showing significant group differences (i.e. SOGS, DSM-V, & KFG; c.f. Table 1), were included as covariates. Response times (RTs) were analysed separately for S1 and S2. The RT model for S1 was set up analogously to the first-stage choice model. The RT model for S2 only included predictors for transition type (common vs. rare) and group (GD vs HC). All main effect predictors (except for group) were entered as within-subject factors to model random effects in all models.

**Table 1.** Demographic Sample Characteristics. GD: gambling disorder group; HC: healthy control group.

|  | GD |  | HC |  |  |  |
| --- | --- | --- | --- | --- | --- | --- |
|  | M | SD | M | SD | t | p |
| N | 30 | - | 30 | - | - | - |
| AGE | 28.63 | 6.37 | 27.27 | 6.15 | 0.85 | 0.4 |
| SCHOOL YEARS | 12.03 | 1.47 | 12.57 | 0.77 | -1.76 | 0.09 |
| MONTHLY INCOME | 1242.07 | 602.91 | 1444.67 | 1643.22 | -0.63 | 0.53 |
| **FTND** | **2** | **2.3** | **0.17** | **0.75** | **4.15** | **2.04\*10<sup>-4</sup>** |
| AUDIT | 6.9 | 7.14 | 7.43 | 4.41 | -0.35 | 0.73 |
| **DSM-V** | **5.47** | **2.32** | **0.2** | **0.41** | **12.27** | **2.17\*10<sup>-13</sup>** |
| **KFG** | **21.1** | **10.11** | **1.93** | **4.88** | **9.35** | **8.36\*10<sup>-12</sup>** |
| **SOGS** | **8.63** | **3.76** | **0.67** | **1.47** | **10.82** | **4.06\*10<sup>-13</sup>** |
| **BDI** | **10.30** | **7.65** | **4.63** | **4.03** | **3.59** | **8.23\*10<sup>-4</sup>** |
| **BIS** | **36.37** | **6.11** | **31.53** | **5.73** | **3.16** | **2.51\*10<sup>-3</sup>** |

#### Model-based analyses Hybrid RL model

We first applied a modified version of the standard hybrid RL model (Daw et al., 2011; Otto et al., 2013) to assess the degree of model-free and model-based control. The model updates MF state-action values (*Q*_*MF*_, Eqs. 1, 2) in both stages via reward prediction errors (RPE; Eqs. 3, 4). In S1, MB state-action values (*Q*_*MB*_) are computed from the transition probabilities and reward estimates using the Bellman Equation (Eq. 5).

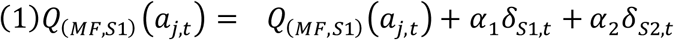

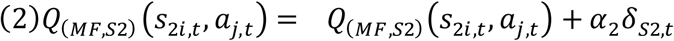

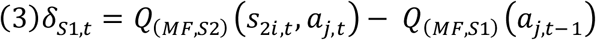

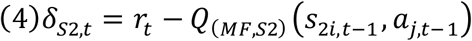

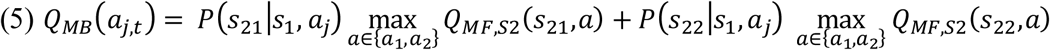

Here, *i* indexes the two different second stages (*S*21, *S*22), *j* indexes actions *a* (*a*_1_, *a*_2_) and *t* indexes trials. Further, *α*_1_ and *α*_2_ denote the learning rates for S1 and S2, respectively. Second - stage MF Q-values (*Q*_(*MF*,*S*2)_) are updated by via reward (*r*_*t*_) prediction errors (*δ*_*S*2,*t*_) (Eq. 2, 4). To model S1 MF Q-values, we allow for reward prediction errors at S2 to influence first-stage Q-values (Eqs. 1, 3).

In addition, Q-values of all unchosen stimuli are assumed to decay (Toyama et al., 2017;2019; Wagner et al., 2022) with a forgetting rate *α*_*decay*_ towards the mean of reward walks (0.5) according to:

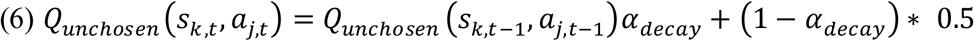

where k ɛ {1,21,22} indexes the three task stages.

#### Softmax Action Selection

In a first step, we used a standard softmax action selection mechanism (Sutton & Barto, 2018) with separate weighting parameters *β* for MF and MB Q-Values in S1 (Eq. 7). The additional parameter *ρ* captures 1st-stage choice perseveration, and weights the indicator function *rep*(*a*) which is set to 1 if the previous S1 choice was the same as the current, and zero otherwise:

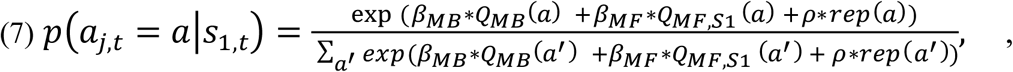

Since S2 is the terminal stage of the task, only MF Q-values factor in (Eq. 8):

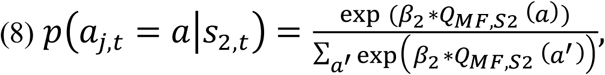

#### Higher-Order Perseveration

In light of the central role of habits and habitual behaviours in theories of addictive disorders, we extended this to the choice perseveration mechanism beyond the trial *t*-1 (Miller et al., 2019; Brands et al. 2025). To this end, the repetition bias *rep*(*a*) in Eq. 7 was replaced with a higher-order perseveration term (HOP, habitual controller, *H*_*i*,*t*_) modelling “value-free” perseveration of familiar options (Miller, 2019). In contrast to the repetition bias *rep*(*a*) in Eq. 7, participant’s entire choice history can exert an influence on current actions. On each trial, *H*_*i*,*t*_ is updated according to:

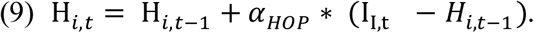

Here, *α*_*HOP*_ is the updating step-size, akin to previously introduced learning and forgetting rates, and models the degree to which the current event (i.e. repetition of choice at t-1) influences *H*_*i*,*t*_. When *α*_*HOP*_=1, the habitual controller H_*i*,*t*_ corresponds to the first-order perseveration mechanism implemented in Eq. 7.

#### Drift Diffusion Action Selection

To more comprehensively examine differences between groups, we replaced softmax action selection with a series of drift diffusion model (DDM)-based choice rules (Pedersen et al., 2017; Shahar et al., 2019; Wagner et al., 2022). In the DDM, choices arise from a noisy evidence accumulation process that terminates as soon as the accumulated evidence exceeds one of two response boundaries. We fitted separate DDMs for each stage of the task, where the upper decision boundary was defined as selection of one stimulus, whereas the lower boundary was defined as selection of the alternative option.

We modelled each stage of the task using separate non-decision time (τ), boundary separation (α) and drift-rate (v) parameters (see Eq. 10). The bias (z) was fixed to 0.5. Response time data were filtered using a percentile-based cut-off, such that on a group-level the fastest and slowest 1% of trials were excluded, and on an individual-subject level, the fastest and slowest 2.5 % were excluded. Single fast outlier trials force the entire RT distribution to adjust to ensure positive density for these observations, which can lead to poor model fit if outlier trials are not taken into account.

We considered three different model variants. The DDM_NULL_ model used fixed drift rates across all trials, i.e. without value modulation. Here, the RT on each trial *t* is distributed according to the Wiener First Passage Time (*wfpt*):

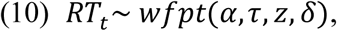

We then examined two models with value-dependent drift-rates (v) either via a linear (DDM_LINEAR_ Eqs. 10, 11) or sigmoid (DDM_SIGMOID_ Eq. 12) transformation of MF and MB Q-values. For the linear version, the S1 drift rate on trial *t* was modelled as:

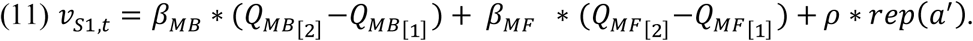

Note that for the HOP-analogue of Equation 10, the indicator function (*rep*(*a*^′^)) is simply swapped out for the habitual controller H_*i*,*t*_ as outlined above (Eq. 9; *β*_*HOP*_ ∗ H_*i*,*t*_).

The DDM_LINEAR_ drift rate in S2 is then:

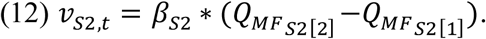

For the non-linear DDM_SIGMOID_, the linear drift rates from Eq. 11 and 12 are additionally passed through a sigmoid function with asymptote 𝑣_*max*_ :

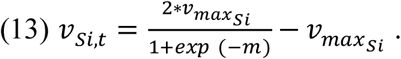

Here, *m* corresponds to the linear drift rate terms from Eq. 11 and 12.

#### Hierarchical Bayesian Model Estimation

Models were fit to all trials and participants, separately for the GD, and HC groups, using a hierarchical Bayesian modelling approach using MCMC sampling as implemented in STAN (Stan Development Team, 2020) running under R (Version 3.5.1) and the *rSTAN* package (Version 2.21.0). Group-level means used normal priors over numerically plausible parameter ranges. Group-level standard deviations used uniform priors over numerically plausible ranges. Priors for all models (DDM and softmax implementations) can be found in the respective model files (https://osf.io/e892f/?view_only=ca199a4ab0ad45f49dafcc464c2e5b66).

Parameters bounded between [0,1] (i.e. learning rates, decay rates, and HOP step-size) were estimated in standard normal space and mapped back onto the interval [0, 1] via the inverse cumulative normal distribution function. Sampling was performed with four chains, each running for 10000 iterations without thinning after a warmup period of 8000 iterations. Chain convergence was assessed via the Gelman-Rubinstein convergence diagnostic *R̂* with 1 ≤ *R̂* ≤ 1.05 for all group and subject-level parameters. Relative model comparison was performed for each group using the *loo-*package in R (Version 2.4.1) via the Widely-Applicable Information Criterion (WAIC) and the estimated log pointwise predictive density (elpd), which estimates the leave-one-out cross-validation predictive accuracy of the model (Vehtari et al., 2017).

Posterior distributions of group differences were assessed using the 95% highest posterior density intervals. We additionally report Bayes Factors for directional effects (dBF) of parameter distributions, estimated via kernel density estimation using R via the RStudio (Version 1.3) interface. These were computed as the ratio of the integral of the posterior difference distribution from 0 to +∞ vs. the integral from 0 to -∞.

### Posterior Predictive Checks

To examine whether the best fitting model reproduced key patterns in the data, we performed posterior predictive checks based on model-derived simulations. To this end, we extracted 500 sets of parameter estimates from the respective model’s posterior distribution and used these to simulate 500 datasets using the *Rwiener* package (Version 1.3.3), and examined the overlap of simulated and observed data for all DDM model variants as a function of value differences. Optimal (i.e. reward-maximising) behaviour was defined separately for each stage as a function of objective value contrast between options (see e.g. Wagner et al., 2022). For each 2^nd^ stage, the objective value difference was defined as the difference in objective reward values (random walk values) between options. For S1, the objective value difference was defined as the difference in model-based values according to Equation 5, with S2 Q-values set their objective maximum. Optimal choices should increase with increasing value differences, and the best-fitting model should reproduce this pattern in both groups.

### Between-Subject Analyses

We tested whether the degree of model-based control in the GD group was associated with clinical measures (addiction severity score, GRCS score, see above) in multiple regression analyses, controlling for covariates of BDI, BIS, FTND and AUDIT. Subject-level mean posterior estimates of *β*_*MB*_ from (1) the softmax model and (2) the DDM_SIGMOID_ served as dependent variables in two separate regression analyses. Due to the superior fit achieved by accounting for HOP, we also included the parameters *β*_*HOP*_ and *α*_*HOP*_ in two exploratory regression analyses.

### FMRI Data Acquisition

MRI images were acquired on a 3 Tesla Magnetom Prisma system (Siemens, Erlangen, Germany) equipped with a 32-channel head coil. Stimuli were presented on an MR compatible screen and mirror system. Participants responded with their index finger on a two-button box, held in their right hand. Functional images were acquired in three separate runs (due to two short breaks, see above) using a multiband (factor 8) gradient echo-planar imaging (mb-EPI) sequence with repetition time (TR) = 0.81 s, echo time (TE) = 30 ms, flip angle = 53°, field of view (FOV) = 212 x 212 mm², voxel size = 2 mm³ isotropic (slice thickness = 2mm). Each volume consisted of 72 transverse slices acquired in alternating order and manually adjusted ∼30° clockwise from the anterior commissure-posterior commissure plane. The three runs contained ∼3000 volumes for each participant with ∼42 min. of total scan time.

### FMRI Data Analysis

#### Preprocessing

Preprocessing and analysis of the fMRI data was performed using SPM12 (v. 7771; Welcome Trust Centre for Neuroimaging, UCL, London, UK) in MATLAB R2024a (Mathworks Inc., Sherborn, MA, USA). Data from each of the three functional runs was preprocessed including realignment, unwarping, and nonlinear normalization to a standard template (Montreal Neurological Institute, MNI) within the DARTEL framework. For normalization purposes, individual high resolution T1-weighted images were used which were co-registered during spatial realignment to the individual mean EPI. Subsequently, normalized images were resliced to a voxel size of 1.5 x 1.5 x1.5 mm^3^ and spatially smoothed with an isotropic 6 mm full-width at half-maximum (FWHM) isotropic Gaussian kernel. A standard high-pass filter with a cut-off at 128s was applied.

#### 1^st^-LevelMmodels

Functional images were analysed using a general linear modelling (GLM) approach as implemented in SPM12. In all GLMs, invalid trials with no responses within the response time window were modelled separately, and all regressors were convolved with the canonical hemodynamic response function as implemented in SPM12. We examined three different 1st - level models to test potential neuronal correlates of model-based and model-free control during task performance.

GLM1 was based on an approach by Kroemer et al. (2019) that mirrors the behavioural analysis of first-stage choice probabilities. First, we included an event regressor for the S1 onset (duration = 0s) with parametric modulators coding for previous relative reward (defined as an outcome greater than the average of the last 20 monetary rewards encountered during the outcome stage), previous transition, and their interaction. Second, the GLM contained separate event regressors (duration = 0s) for the S2 onset following a common (70%) or rare (30%) transition. Onset-related BOLD-activation for both trial types was allowed to vary according to both model-free and model-based PEs (Daw et al., 2011). While MF PEs at S2 compare MF Q-values of 1st- and 2nd-stage stimuli, MB-PEs reflect the difference between first-stage MB Q-values and second-stage MF- action-values. Lastly, GLM1 included an event regressor (duration = 0s) for the outcome onset, parametrically modulated by the MF PEs (difference between outcome and MF Q-value).

GLM2 was constructed similarly, with two key differences. First, we collapsed rare and common events at S2, and included two parametric modulators (one for MF-PE and one for MB-PE) for this event to increase robustness in the estimation of the value-related BOLD response. Second, we included two separate categorical outcome regressors, according to relative reward magnitude, by subtracting the mean of the last twenty trials from the reward received on a given trial, and classifying trials as either high or low relative reward magnitude.

GLM3 explored whether differences in model-based behaviour were reflected in differential S1onset effects. We constructed four S1 onset regressors (duration = 0s) based on previous reward (high vs. low magnitude) and transition (rare vs. common). This resulted in four event types: high reward + common transition, high reward + rare transition, low reward + common transition, and low reward + rare transition. We reasoned differences in task structure learning and/or integration of transition probabilities into choice might manifest in distinct first-stage onset effects.

#### *2^nd^-Level* Models

All second-level fMRI models included covariates of depressive symptoms (BDI-II), nicotine use (FTND), alcohol use (AUDIT) and age.

Basic group comparisons used two-sample t-tests at the second level, with group (GD vs. HC) as a between-subject factor, focusing on the following first-level contrasts: (1) first-stage parametric contrast images of previous reward magnitude and previous transition, and (2) parametric contrast images of MF RPE at outcome onset. Flexible factorial models with the factors subject, condition and group, were used to assess group differences between rare and common S2 onsets (GLM 1), MF- and MB-PEs (GLM 2), high vs. low reward outcomes (GLM 2) and S1 onset effects as a function of reward and transition (GLM 3).

Whole-brain effects were corrected for multiple comparisons using FDR-correction (peak-level; p<.05). Effects related to MF/MB PEs and reward used correction for multiple comparisons using an *a priori* defined region of interest (ROIs) and small-volume-correction (SVC, p<0.05, FWE correction). We used a combined mask based on two meta-analyses, provided by the Rangel Neuroeconomics lab (http://www.rnl.caltech.edu/resources/index.html). This mask covers bilateral ventral striatum, vmPFC, posterior cingulate cortex (PCC), and anterior cingulate cortex (ACC).

#### Exploratory Analysis: Model-Based Control and Neural Prospection Signatures

In a final exploratory analysis, we investigated the association between model-based control (*β*_*MB*_) and neuronal correlates of vivid prospection. This was motivated by the idea that model-based control may rely on the mental simulation of future outcomes, similar to prospection. Reduced model-based control in GD my then be due to a reduction in vivid prospection, whereas higher model-based control may be associated with more vivid mental imagery of future outcomes. To this end, we applied a recently-developed whole-brain decoder for prospection vividness (Lee et al., 2022) based on a thresholded partial least squares algorithm (Lee, Bradlow, & Kable, 2022) that predicts the vividness of prospection from whole-brain maps. This decoder was applied to the S1 onset contrast images for each participant, yielding a neural vividness score for each participant.

In short, the decoder was constructed based on 24 participants that imagined various future scenarios for 12 seconds each. These scenarios were selected using a 2 × 2 design that varied independently in vividness (vivid vs. non-vivid) and valence (positive vs. negative). Using a leave-one-out cross-validation approach, Lee et al. (2022) demonstrated that the neural predictor for vividness could reliably distinguish vivid from non-vivid scenarios but did not predict valence. Conversely, the valence predictor successfully classified positive and negative scenarios but was unrelated to vividness.

To apply the decoder to our data, we first down-sampled the stage 1 onset contrast images from a 1.5mm template to a 2mm MNI standard template to match the dimensions of the vividness predictor. We then computed a voxel-wise dot product between the contrast images and the neural predictor, yielding a single "vividness readout" for each participant. While the absolute scale of this readout depends on preprocessing factors such as grand mean scaling, higher values indicate a greater likelihood, based on the decoder model, that a participant engaged in vivid imagery / prospection.

## Results

### Sample Characteristics

Groups were matched on age, school years and monthly income (see Table 1). As expected, the GD group exhibited higher levels of problematic gambling compared to the HC group (DSM-V, KFG scores, SOGS scores, see Table 1). The GD group furthermore exhibited significantly higher levels of nicotine use (FTND), depressive symptoms (BDI-II) and trait impulsivity (BIS-15.).

### Model-agnostic behavioural results

In line with previous work, the GD group showed reduced task performance, as reflected in significantly lower average rewards earned per trial throughout the task (M ± SE: GD (HC) 60.07 (62.15) ± 0.71 (0.41); t=-2.36, p=0.02; Figure 1A). Following standard analysis approaches of TST-behaviour we examined the stay probability *p*(stay), i.e. the tendency to repeat the previous S1 action as a function of outcome reward magnitude and transition. MB behaviour is based on both outcome (reward) and transition (utilizing a task model), whereas MF behaviour is solely based on previous reinforcement (e.g. Daw et al., 2011; Otto et al., 2013). The linear mixed model predicting *p*(stay) included transition (common vs. rare), reward (high vs. low), and group (HC vs. GD) along with their interactions as predictors. We observed significant main effects of group (β=-0.42, p<.001) and reward (β=0.10, p<.001; indicating a MF contribution, Figure 3), and a significant reward*transition interaction (β=0.38, p<.001) indicating MB control (Supplementary Table S2; Figure 1 D-E). In addition, the three-way interaction was significant (β=-0.10, p=.027), reflecting a reduced MB effect in the GD group (see Figure 1D, 1E). None of the clinical covariates showed significant effects (Supplementary Table S2).

**Figure 1.**
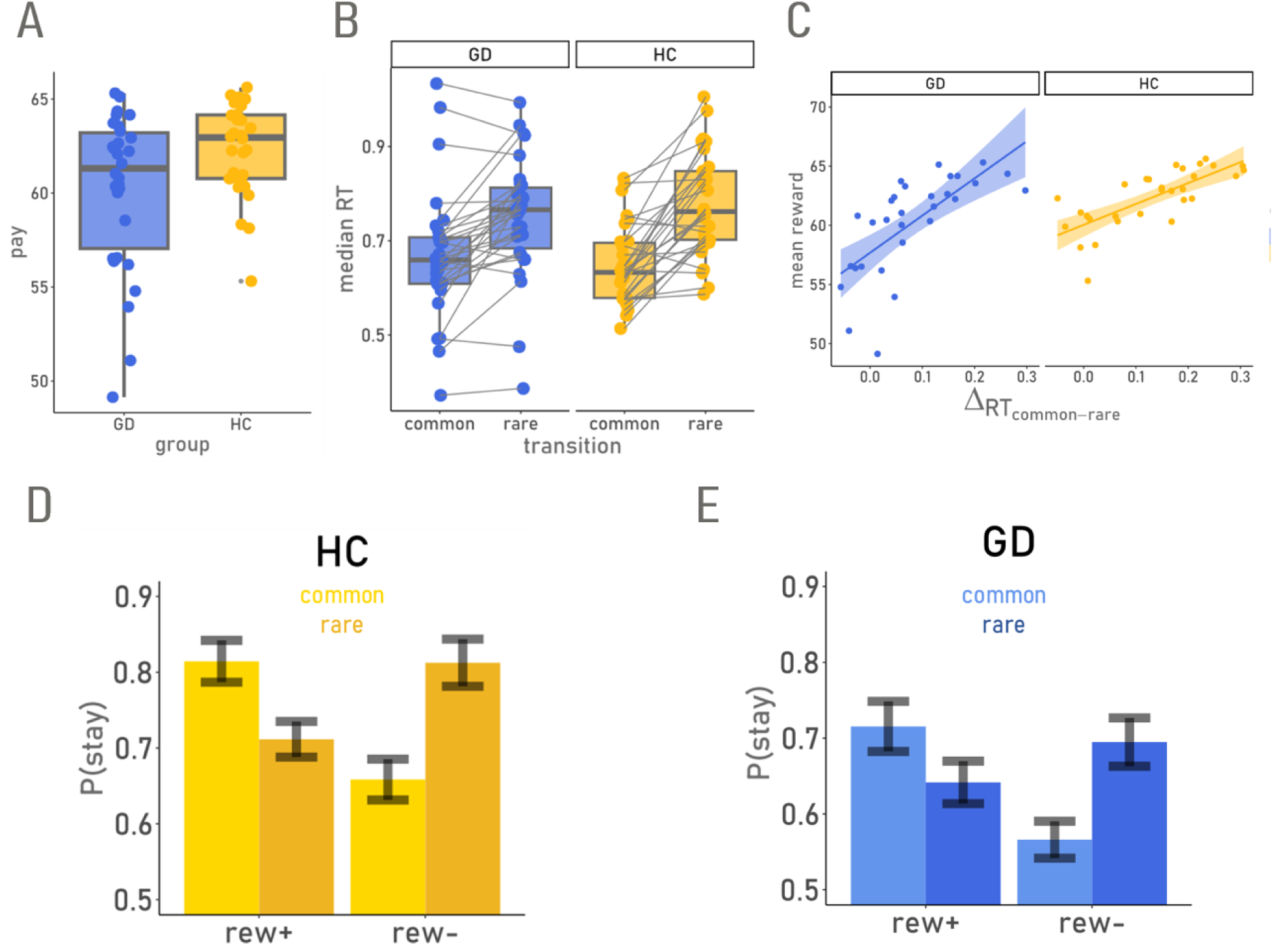
Model-agnostic behavioral data (blue: gambling disorder [GD] group, yellow: healthy control [HC] group). Points represent data from individual participants. A: Mean rewards earned per trial. B: Median reaction times at the second stage as a function of preceding transition. C: Association of RTs following common vs. rare transitions (delta RT) and mean rewards earned. D-E: Stay probabilities from the HC (D) and GD group (E). Y-axis: probability of repeating the S1 choice of the previous trial (p(stay)). Lighter/darker shades depict common/rare transitions. Rew+/- indicates dichotomisation of previous trials into high (rew+) and low reward trials (rew-) compared to the average reward obtained in the preceding 20 trials.

Another index of MB control is the modulation of S2 RTs by transition type. Specifically, in participants that have an accurate model of the task transition structure, rare transitions should elicit longer S2 RTs, reflecting unexpected events. Both groups showed longer S2 RTs following rare transitions (main effect transition: β=-0.001; p<.001; Figure 1B, Supplementary Table S3), but the effect was significantly more pronounced in the HC vs. GD group (transition*group interaction in the LMM predicting S2 RTs, β=0.003, p=.016; Supplementary Table S3). Consistent with the idea that MB behaviour is overall beneficial for task performance, this RT-difference was significantly correlated with the average payout in both groups (HC: R=0.77, p<.001; GD: R=0.80, p<.001; Figure 1B, 1C).

### Model-Based Analysis

#### Model Comparison

Model comparison for the softmax choice rule models revealed that the hybrid model accounting for HOP provided a superior fit in both groups (Table 5). Therefore, this model variant was used for the DDMs with value-modulation of drift rates (i.e. DDM_LINEAR_ and DDM_SIGMOID_). Model comparison across all three DDMs (DDM_NULL_, DDM_LINEAR,_ DDM_SIGMOID_) are provided in Table 6. We observed the same model ranking in both groups, such that the DDM_SIGMOID_ provided the best account of the data.

**Table 5.** Model comparison of softmax models.

| GD |  |  |  | HC |  |  |
| --- | --- | --- | --- | --- | --- | --- |
| model | eldp<sub>diff</sub> | se<sub>diff</sub> | rank | eldp<sub>diff</sub> | se<sub>diff</sub> | rank |
| Q + HOP | 0.0 | 0.0 | 1 | 0.0 | 0.0 | 1 |
| Q + FOP | -34.5 | 15.5 | 2 | -27.1 | 10.9 | 2 |

Note. GD: Gambling Disorder group. HC: Healthy Control group. FOP/HOP: First-Order Perseveration/Higher-Order Perseveration. eldp<sub>diff</sub>: difference in estimated log pointwise predictive density.

**Table 6.** Model comparison of reinforcement learning drift diffusion models (RLDDMs)

| GD |  |  |  | HC |  |  |
| --- | --- | --- | --- | --- | --- | --- |
| model | eldp<sub>diff</sub> | se<sub>diff</sub> | rank | eldp<sub>diff</sub> | se<sub>diff</sub> | rank |
| DDM<sub>SIGMOID</sub> | 0.0 | 0.0 | 1 | 0.0 | 0.0 | 1 |
| DDM<sub>LINEAR</sub> | -345.4 | 57.5 | 2 | -360.2 | 61.6 | 2 |
| DDM<sub>NULL</sub> | -3112.2 | 352.5 | 3 | -4383.2 | 388.1 | 3 |

Note. GD: Gambling Disorder group. HC: Healthy Control group. null/linear/sigmoid: DDM versions without/with linear/with sigmoid drift rate scaling. eldp<sub>diff</sub>: difference in estimated log pointwise predictive density.

### Posterior Predictive Checks

To verify that the best-fitting model could reproduce key patterns in the data, we simulated and analysed 500 full data sets from the posterior distribution of the three DDMs. The proportion of correct choice predictions per model variant and task stage aligned with the previously shown model ranking (see Supplementary Table S4). Additionally, we examined choice behaviour as a function of choice difficulty. For both S1 and S2, we focused on optimal choices based on Q-value differences (Figure 2, see methods). Again, the DDM_SIGMOID_ accurately reproduced changes in performance as a function of Q-value differences, both for S1 (Figure 2A, 2B) and S2 (Figure 2C, 2D), and in both groups.

**Figure 2.**
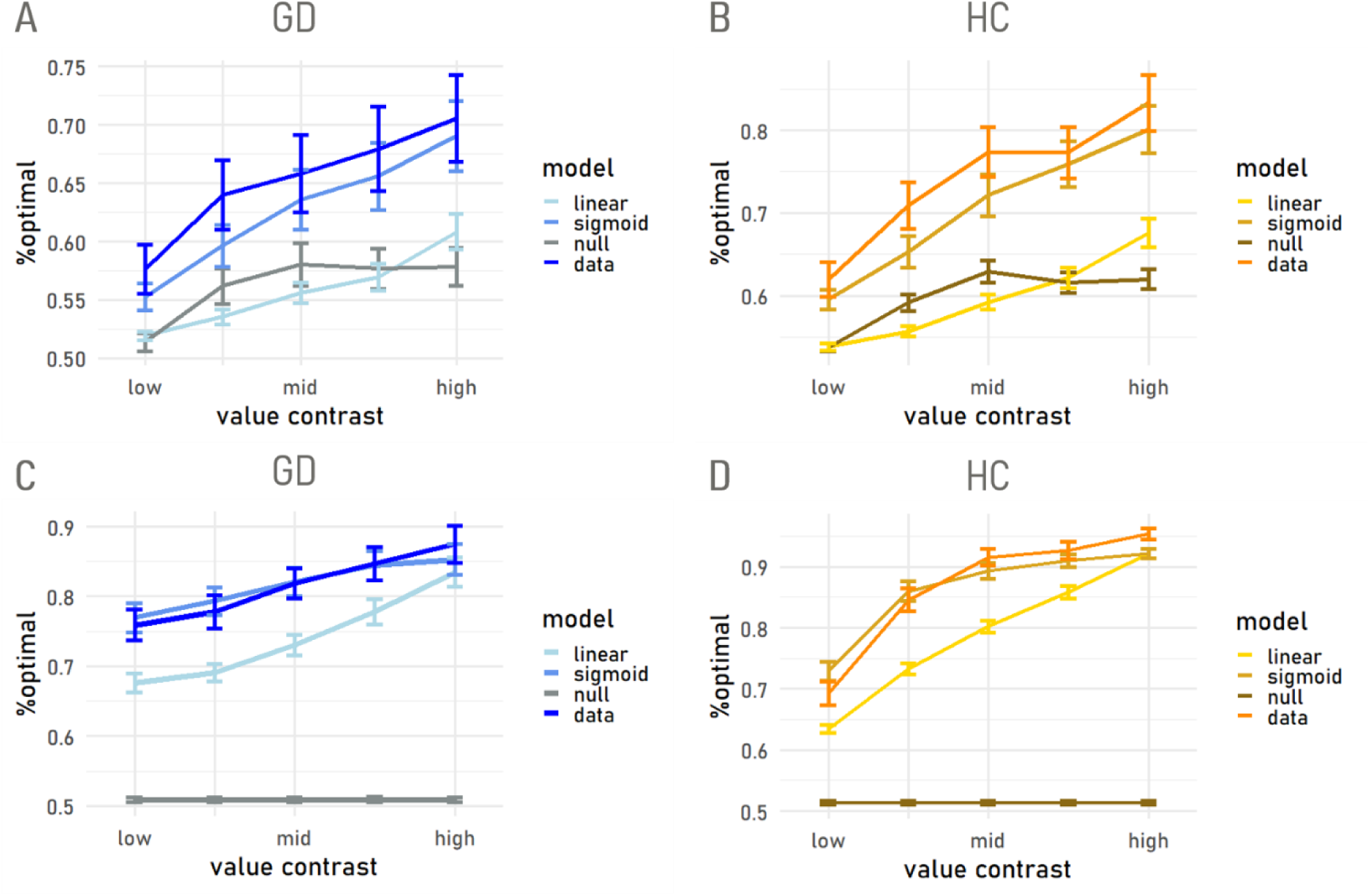
Posterior predicitive checks for reinforcement learning drift diffusion models (DDMS) for stage 1 (A, B) and stage 2 (C, D). Null model: DDM without value modulation. Linear: DDM with linear drift rate modulation. Sigmoid: DDM with non-linear drift rate modulation. Data: observed choice behaviour. Value contrast (x-axis): binned Q-value differences of S1 options (A, B) / binned reward differences of S2 options (C, D). Y-axis plots the proportion of optimal observed / model-predicted choices. Error bars depict the standard error across participants (n=30) and simulations (n=500).

### Parameter Estimates

Posterior parameter estimates from the hybrid RL model (QL+HOP) mirrored the reduced MB control in the GD group observed in the model-agnostic analyses above. Model-based control was reduced in the GD group (95% HDI *β*_*MB*−*DIFF*_ : [-11.81, -0.43], dBF(< 0) = 75.38, Table 7, c.f. Table 4, Figure 3A). The HOP-stepsize tended to be lower in the GD group (95% HDI *α*_*HOP*−*DIFF*_ : [-0.78, 0.03], dBF(< 0) = 21.60, Table 7; Figure 3B), although the 95% HDI of the difference distribution showed minimal overlap with 0.

**Figure 3.**
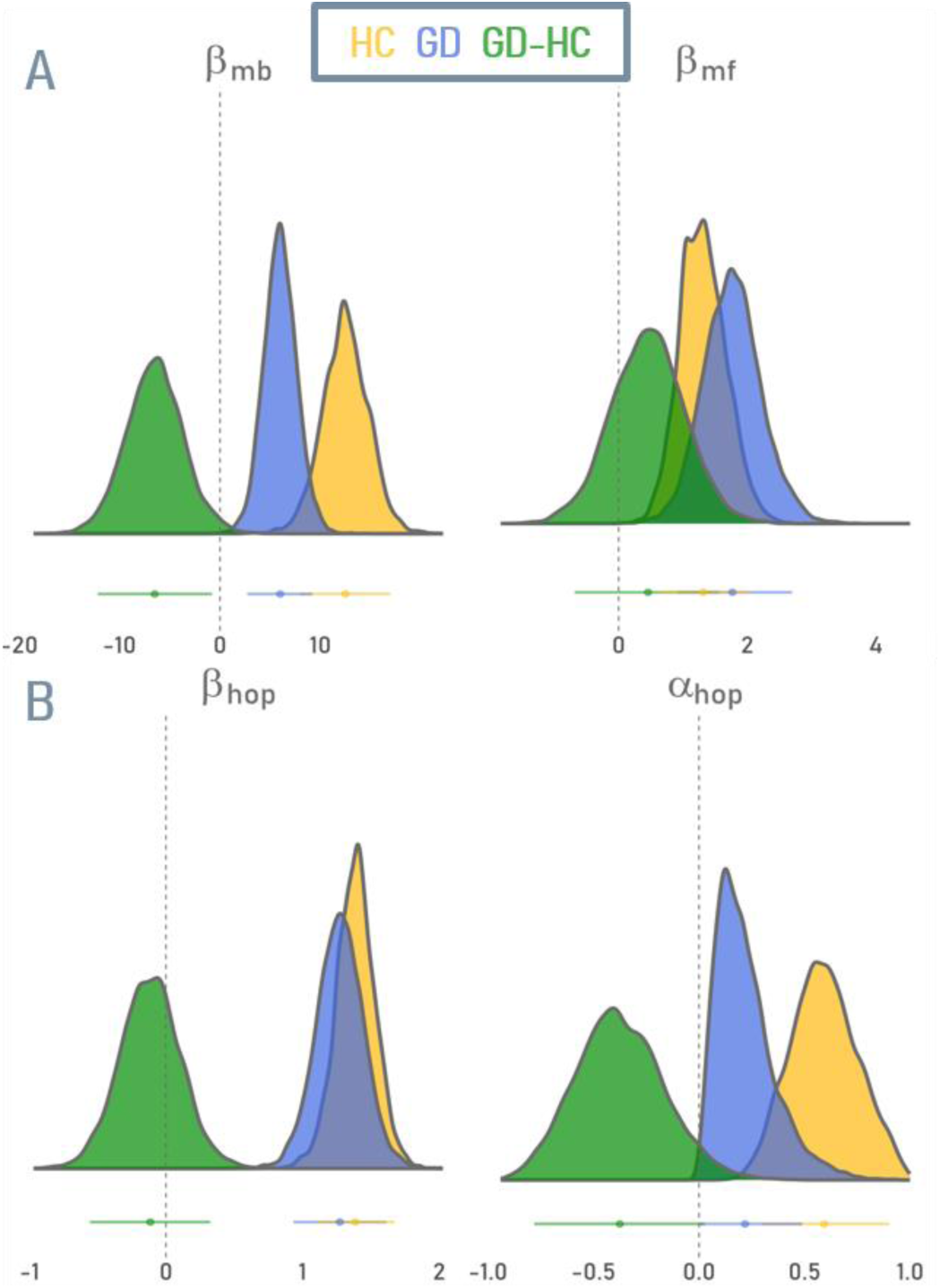
Posterior distributions of selected parameters from the best-fitting softmax hybrid RL model with higher-order perseveration. A: model-based (*β*_*MB*_) and model-free (*β*_*MF*_) action control,. B: choice weight (*β*_*HOP*_) and stepsize (*α*_*HOP*_) parameters for the habitual controller (HOP). Yellow: control group (HC), blue: gambling group (GD); green: posterior difference distribution (GD – HC).

**Table 7.**
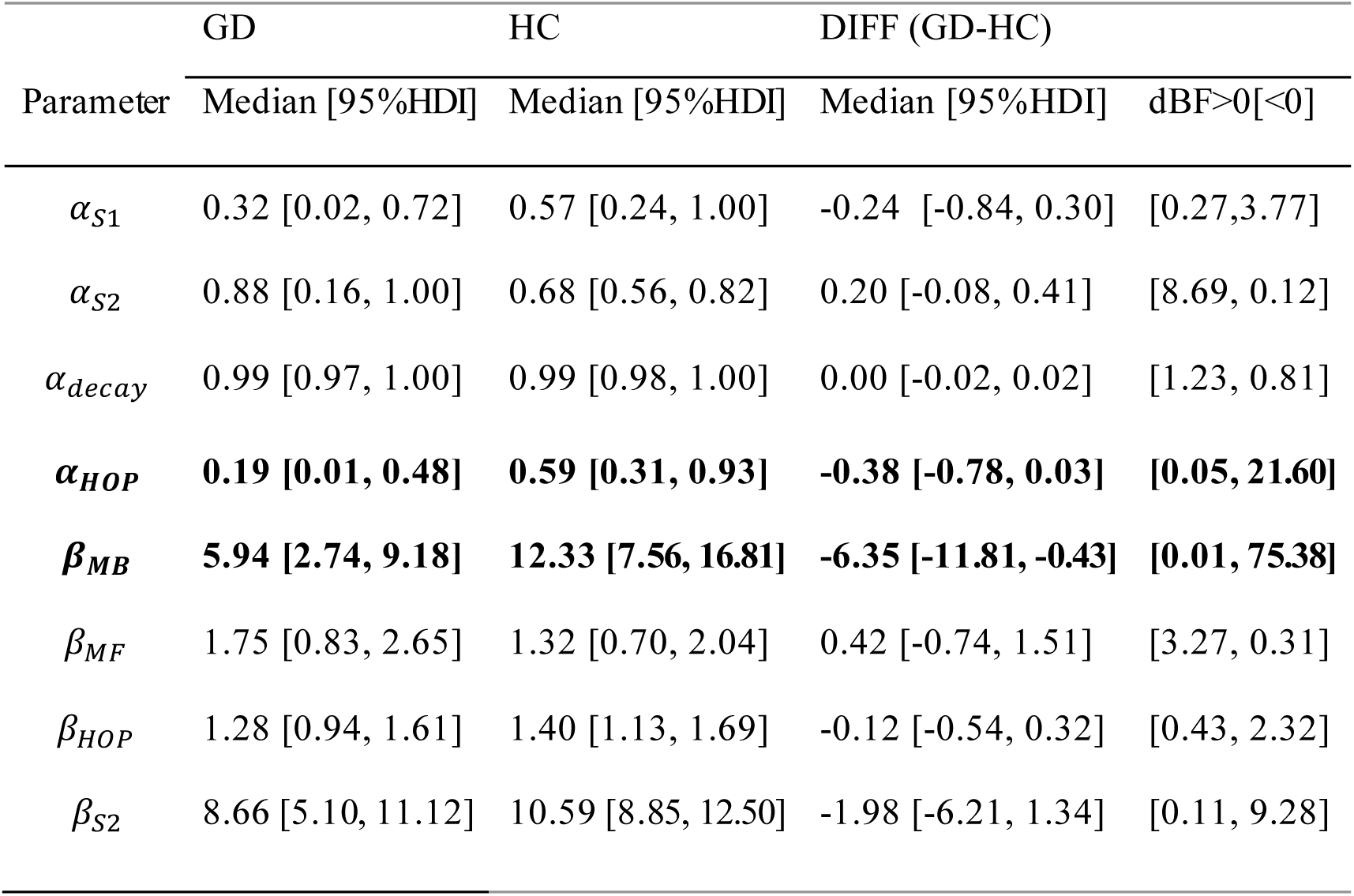
Group mean posterior parameter estimates from the best-fitting softmax model with HOP.

|  | GD | HC | DIFF (GD-HC) |  |
| --- | --- | --- | --- | --- |
| Parameter | Median [95%HDI] | Median [95%HDI] | Median [95%HDI] | dBF>0[<0] |
| $\alpha_{S1}$ | 0.32 [0.02, 0.72] | 0.57 [0.24, 1.00] | -0.24 [-0.84, 0.30] | [0.27,3.77] |
| $\alpha_{S2}$ | 0.88 [0.16, 1.00] | 0.68 [0.56, 0.82] | 0.20 [-0.08, 0.41] | [8.69, 0.12] |
| $\alpha_{decay}$ | 0.99 [0.97, 1.00] | 0.99 [0.98, 1.00] | 0.00 [-0.02, 0.02] | [1.23, 0.81] |
| $\alpha_{HOP}$ | **0.19 [0.01, 0.48]** | **0.59 [0.31, 0.93]** | **-0.38 [-0.78, 0.03]** | **[0.05, 21.60]** |
| $\beta_{MB}$ | **5.94 [2.74, 9.18]** | **12.33 [7.56, 16.81]** | **-6.35 [-11.81, -0.43]** | **[0.01, 75.38]** |
| $\beta_{MF}$ | 1.75 [0.83, 2.65] | 1.32 [0.70, 2.04] | 0.42 [-0.74, 1.51] | [3.27, 0.31] |
| $\beta_{HOP}$ | 1.28 [0.94, 1.61] | 1.40 [1.13, 1.69] | -0.12 [-0.54, 0.32] | [0.43, 2.32] |
| $\beta_{S2}$ | 8.66 [5.10, 11.12] | 10.59 [8.85, 12.50] | -1.98 [-6.21, 1.34] | [0.11, 9.28] |

**Table 9.** Group mean posterior parameterestimatesfrom the best-fitting RLDDM (DDM_SIGMOID_) with HOP.

|  | GD | HC | DIFF (GD-HC) |  |
| --- | --- | --- | --- | --- |
| parameter | Median [95%HDI] | Median [95%HDI] | Median [95%HDI] | dBF>0[<0] |
| $\alpha_{S1}$ | 1.24 [1.17, 1.31] | 1.26 [1.18, 1.34] | -0.03 [-0.12, 0.08] | 0.43 [2.30] |
| $\alpha_{S2}$ | 1.44 [1.36, 1.51] | 1.38 [1.33, 1.43] | 0.06 [-0.03, 0.15] | 8.02 [0.12] |
| $\tau_{S1}$ | **0.31 [0.27, 0.34]** | **0.35 [0.33, 0.38]** | **-0.04 [-0.09, 0.00]** | **0.02 [44.05]** |
| $\tau_{S2}$ | **0.33 [0.29, 0.37]** | **0.38 [0.36, 0.40]** | **-0.05 [-0.10, -0.01]** | **0.01 [169.00]** |
| $\nu_{MAX-S1}$ | **0.86 [0.04, 1.52]** | **1.81 [1.27, 2.32]** | **-0.93 [-1.89, 0.01]** | **0.03 [31.82]** |
| $\nu_{MAX-S2}$ | **1.68 [1.45, 1.92]** | **1.96 [1.82, 2.11]** | **-0.28 [-0.55, 0.00]** | **0.02 [40.13]** |
| $\nu_{MB}$ | 11.72 [6.88, 16.69] | 17.42 [12.36, 20.21] | -5.71 [-12.71, 1.57] | 0.07 [13.69] |
| $\nu_{MF}$ | 3.29 [2.17, 4.53] | 2.00 [0.93, 3.22] | 1.30 [-0.40, 2.97] | 14.33 [0.07] |
| $\nu_{S2}$ | 16.47 [12.15, 21.04] | 16.21 [12.52, 20.32] | 0.34 [-5.28, 6.30] | 1.21 [0.82] |

Posterior parameter estimates from the best-fitting DDM model (DDM_SIGMOID_) showed similar, though less pronounced differences in MB control between groups, (95% HDI 𝑣_*MB*−*DIFF*_ : [- 12.71, 1.57], dBF(𝒗_***MB***− ***DIFF***_) = 13.96; M=-5.71; Table 8). For both stages, the GD group exhibited considerably lower non-decision times compared to the HC group (95% HDI *τ*_*S*1−*DIFF*_ : [-0.09, 0.00], dBF(< 0) = 44.05; 95% HDI *τ*_*S*2−*DIFF*_ : [-0.10, -0.01], dBF(< 0) = 169.0; c.f. Table 8; Figure 4A, 4B). This pattern was also mirrored in drift rate asymptotes at both stages, such that the HC group had lower maximum drift rates in S1 (95% HDI *ν*_*MAX*−*S*1−*DIFF*_: [-1.89, -0.01], dBF(< 0) = 31.82) and S2 (95%HDI *ν*_*MAX*−*S*2−*DIFF*_: [-0.55, 0.00], dBF(< 0)= 40.13).

### Correlation Analyses

When examining possible associations of MB control (*β*_*MB*_, 𝑣_*MB*_), with gambling severity, gambling-related cognitions (GRCS score) or clinical covariates in the GD group, no significant effects were observed (all ps>.1; Supplementary Table S5). For HOP parameters from the softmax and DDM_SIGMOID_, *β*_*HOP*_ was associated with impulsivity (BIS) (ß= -0.03; p= .006). The step size parameter (α_*HOP*_) was significantly associated with cognitive distortions in the GD group (softmax: ß=-0.03, p=.03; RLDDM: ß=-0.03, p=.02; Supplementary Table S6), such that higher GRCS scores were linked to a lower HOP-step size (i.e. an extended window of habitual response integration).

Based on attenuated effects of MB control in the DDM_SIGMOID_ compared to the softmax model, we next examined potential parameter trade-offs in the DDM implementation. As can be seen in Figure 6, *β*_*MB*_ from the softmax model was significantly correlated with S1 non-decision times and S1 maximum drift rates from the DDM_SIGMOID_, suggesting that the MB control deficits in the GD group were partly accounted for by changes in these other DDM_SIGMOID_ components, beyond *ν*_*MB*_.

**Figure 5.**
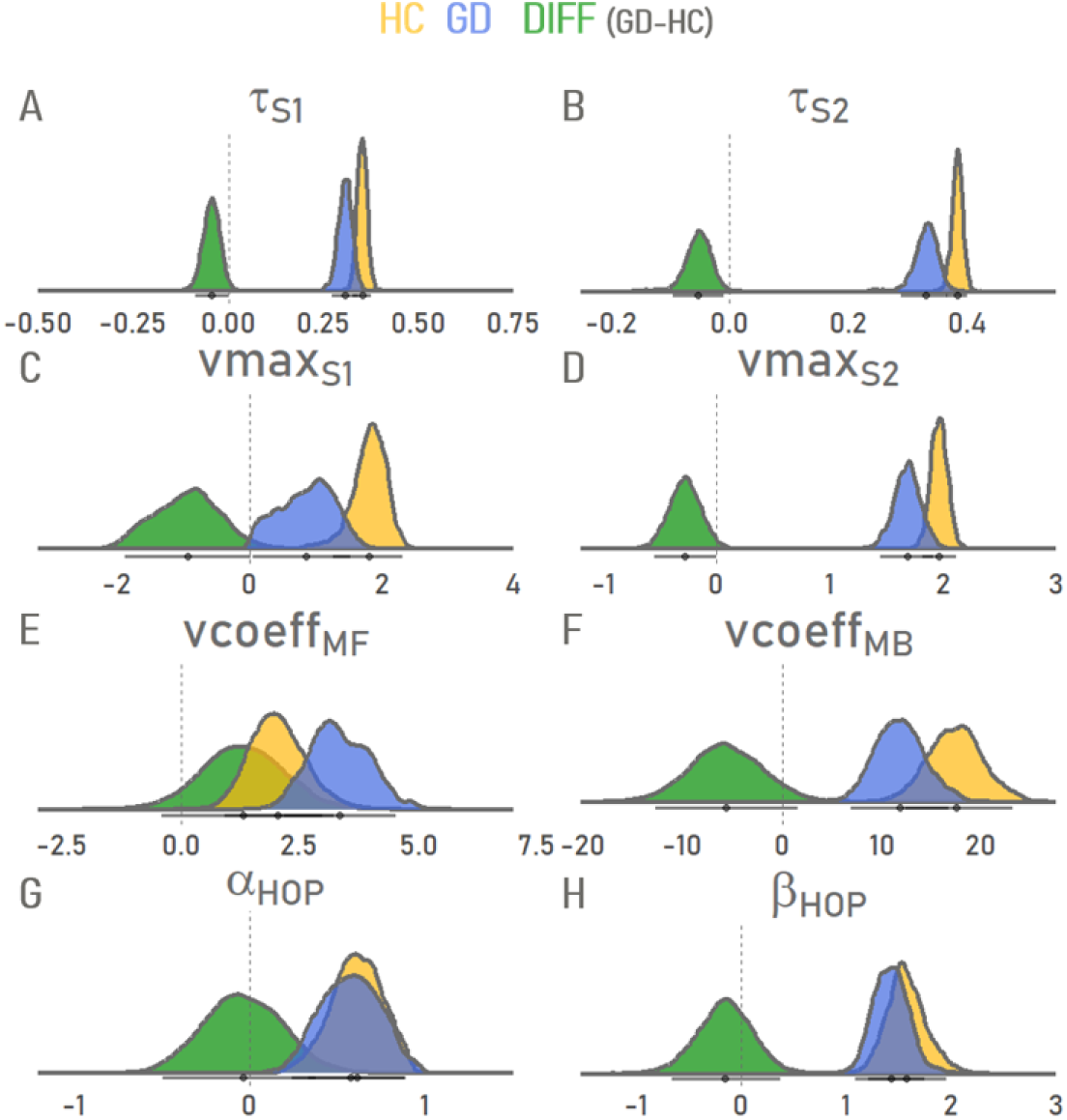
Posterior distributions of selected parameters from the best-fitting RLDDM with nopn-linear drift rate scaling. A-B: non-decision time parameters for S1 and S2. C-D: drift rate asymptotes for S1 and S2. E-F: model-free and model-based drift rate coefficients. G-H: step-size and choice weight parameters for the habitual controller. Yellow: control group (HC), blue: gambling group (GD); green: posterior difference distribution (GD – HC).

**Figure 6.**
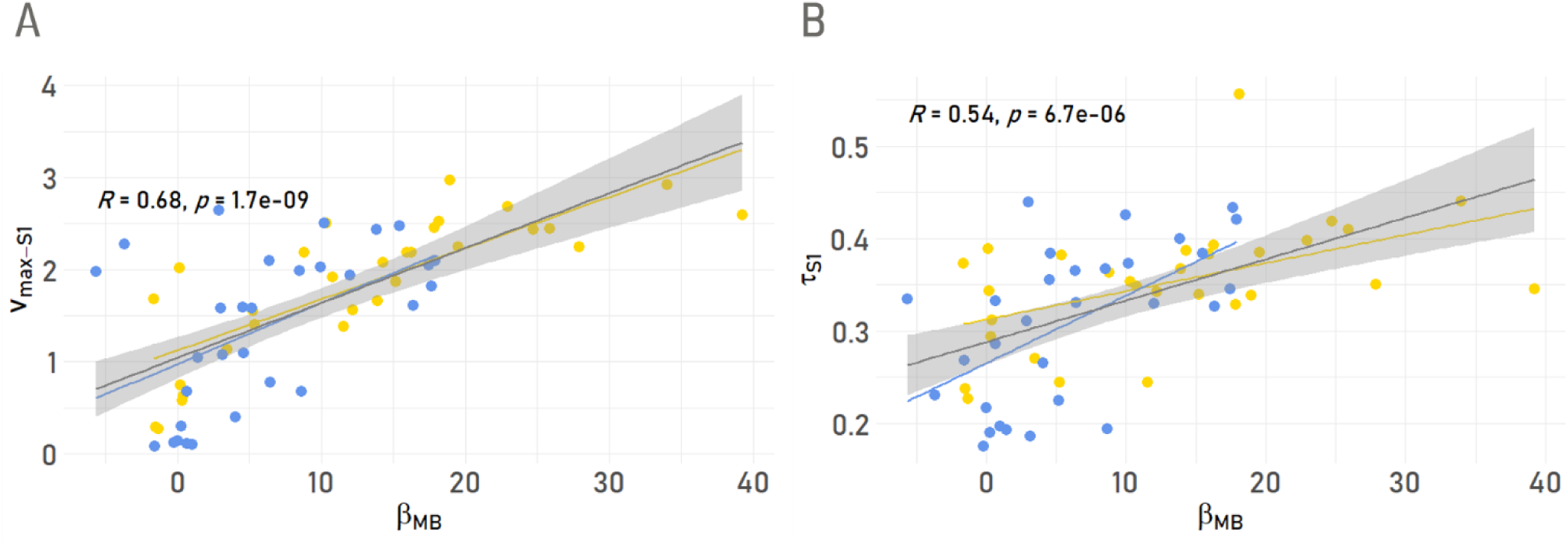
Associations between S1 RLDDM parameters and model-based control from the softmax model. ß_MB_: Softmax-model derived parameter for MB control. A: association with S1 maximum drift rate/ asymptote. B: association with S1 non-decision time. Blue/yellow: data from participants in the GD/HC group, respectively.

### FMRI Results

#### S1 Onset Effects (GLM1)

We first examined BOLD activity at the onset of the first-stage, using an event-related regressor that was modulated by previous reward and transition (Kroemer et al., 2019) (see methods), to investigate potential mechanisms underlying task structure updating. The contrast of previous reward (high > low, Figure 7) revealed widespread cortical and subcortical effects, including prominent bilateral hippocampal clusters (left: x/y/z = -28.5/-9/-16.5); z-value = 6.26; p<0.001; right: x/y/z = 30/-10.5/-19.5); z-value = 5.21; *p* < 0.001) on the whole brain level. No significant group differences were observed.

**Figure 7.**
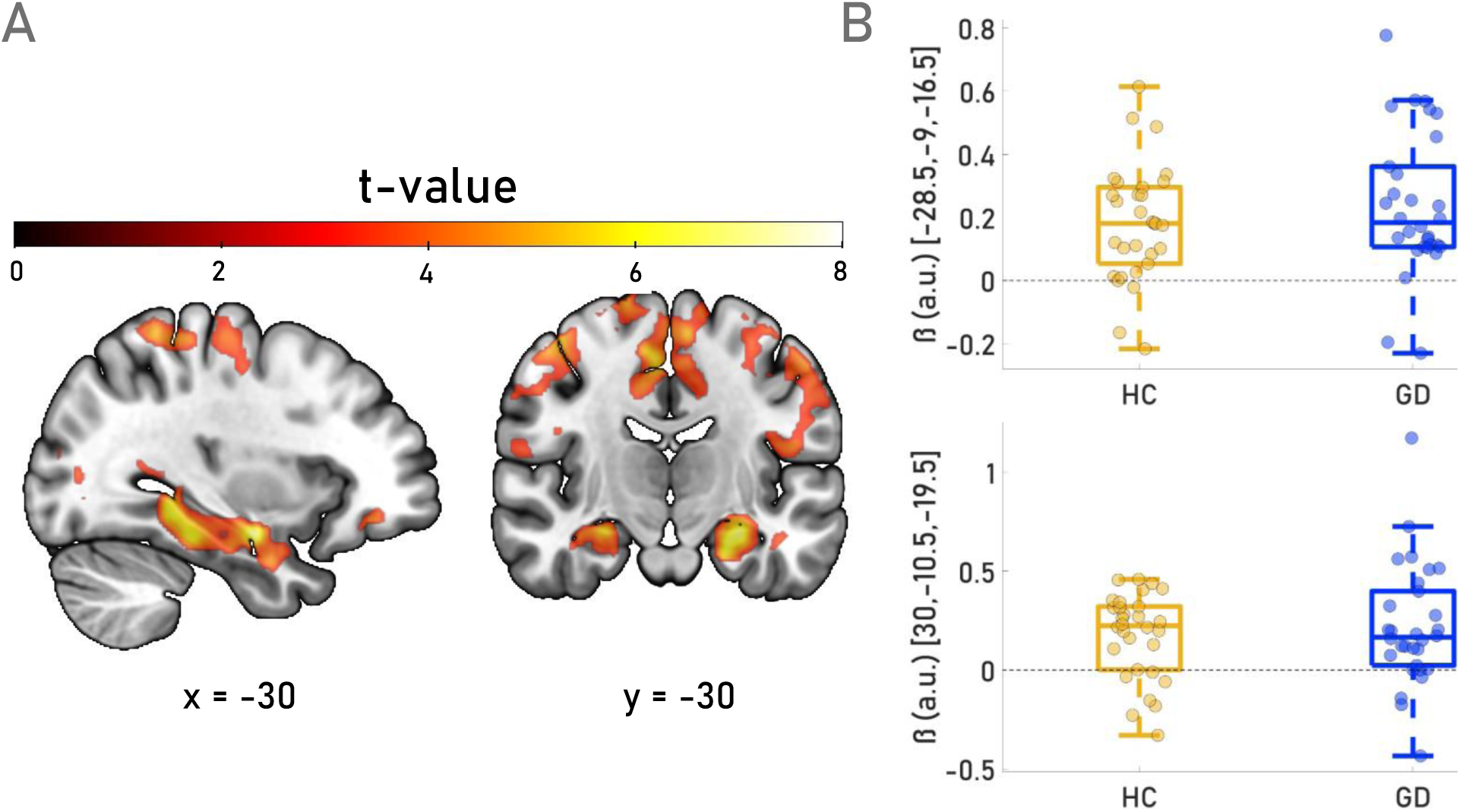
Main effect of previous-trial reward on BOLD activity at first-stage onset. (**A**) Whole-brain statistical parametric map illustrating the main effect of previous-trial reward on BOLD signal at S1 onset (parametric modulation analysis; *p* < 0.001 uncorrected). Prominent bilateral activation is observed in medial temporal lobe regions, including the hippocampus and basolateral part of the amygdala (BLA). (**B**) Parameter estimates (beta weights) extracted from peak voxels in the left [-28.5, -9, -16.5] and right [30, -10.5, -19.5] medial temporal lobe clusters, corresponding to regions of maximal activation in (A); a.u. = arbitrary units. Boxplots depict group -level distributions for GD (blue) and HC (yellow). No significant group differences were observed at either site.

The GLM1 contrast of previous trial transition (rare > common) then probed neural sensitivity to task structure and potential model-based updating signals, and revealed engagement of prefrontal regions, with a prominent cluster centered in the anterior medial prefrontal cortex (amPFC; x/y/z = -7.5/58.5/10.5); z-value = 5,16; *p* = 0.022 (whole brain corrected)). However, again, no reliable group differences were observed (Figure 8).

**Figure 8.**
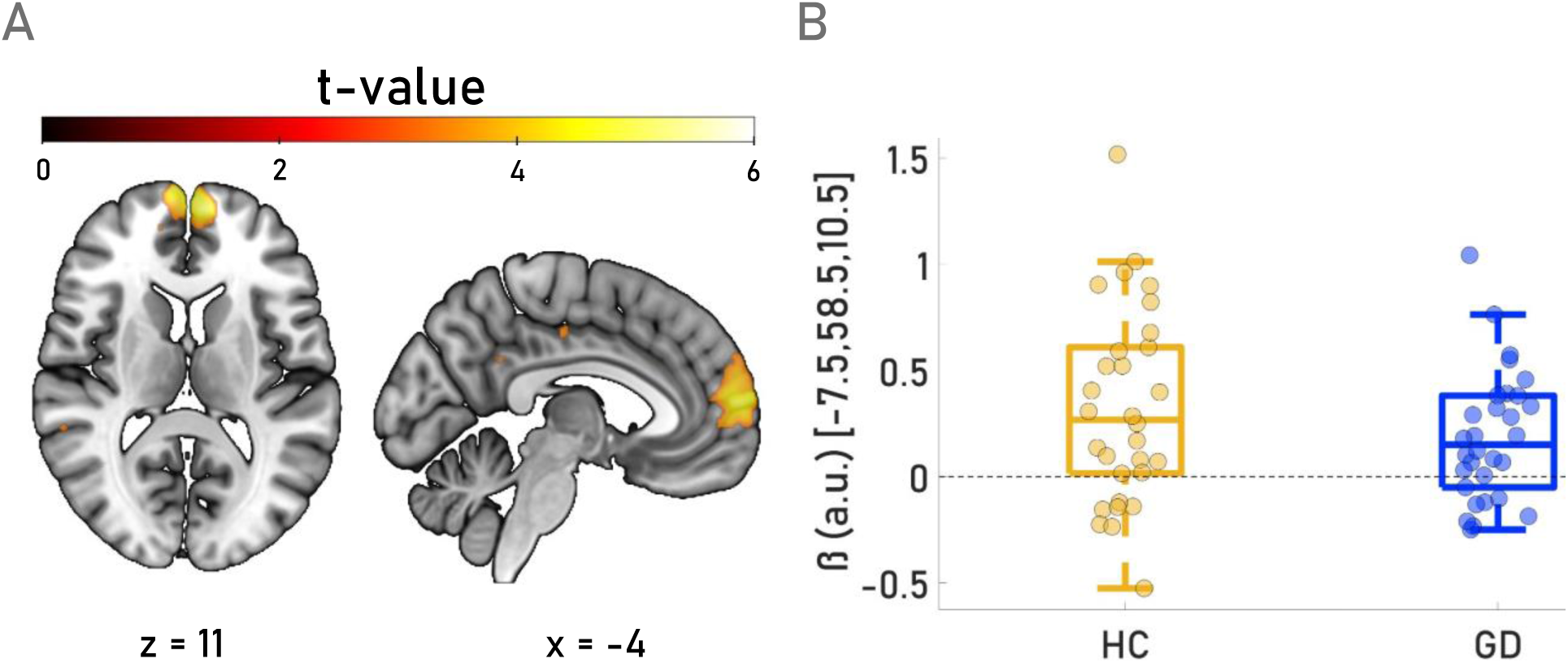
Neural response to previous-trial transition type (rare > common) at first-stage onset. **(A)** Whole-brain activation map displaying the main effect of previous-trial transition type (parametric modulation at S1 onset; *p* < 0.001 uncorrected). Robust activation was observed in the medial prefrontal cortex, with a peak located at MNI coordinates [-7.5, 58.5, 10.5], corresponding to the anterior medial prefrontal cortex (amPFC). **(B)** Beta estimates extracted from the peak voxel in the amPFC. A.u. = arbitrary units; Boxplots depict the distribution of parameter estimates for GD (blue) and HC (yellow). No significant group differences were observed.

Similarly, we did not find any significant group differences for the contrast of previous reward x previous transition.

#### S2 Onset Effects (GLM1)

We next examined transition effects on S2 onset-related activation. Increased engagement of e.g. executive control regions following rare vs. common transitions would be expected if participants correctly understood the task transition structure of the task (similar to previously reported state prediction errors, Gläscher et al. 2010). We hypothesized that group differences in these effects might underlie group differences in model-based behavior. Rare vs. common transitions (on the whole-brain level) indeed elicited stronger activation in a distributed frontoparietal network previously implicated in state prediction errors (left inferior frontal gyrus: x/y/z = -32/26/4), z-value = 6.03; p < 0.001; dorsomedial prefrontal cortex: x/y/z = 2/16/51, z-value = 5.91, p < 0.001), right inferior parietal lobule: x/y/z = 34/-51/44], z-value = 5.01, p < 0.001) – all p-values are family wise error (FWE) corrected - (see e.g. Alexander & Brown, 2011; Silvetti et al., 2014) (see Figure 9). However, no significant interaction effects with group were observed at these peaks.

**Figure 9.**
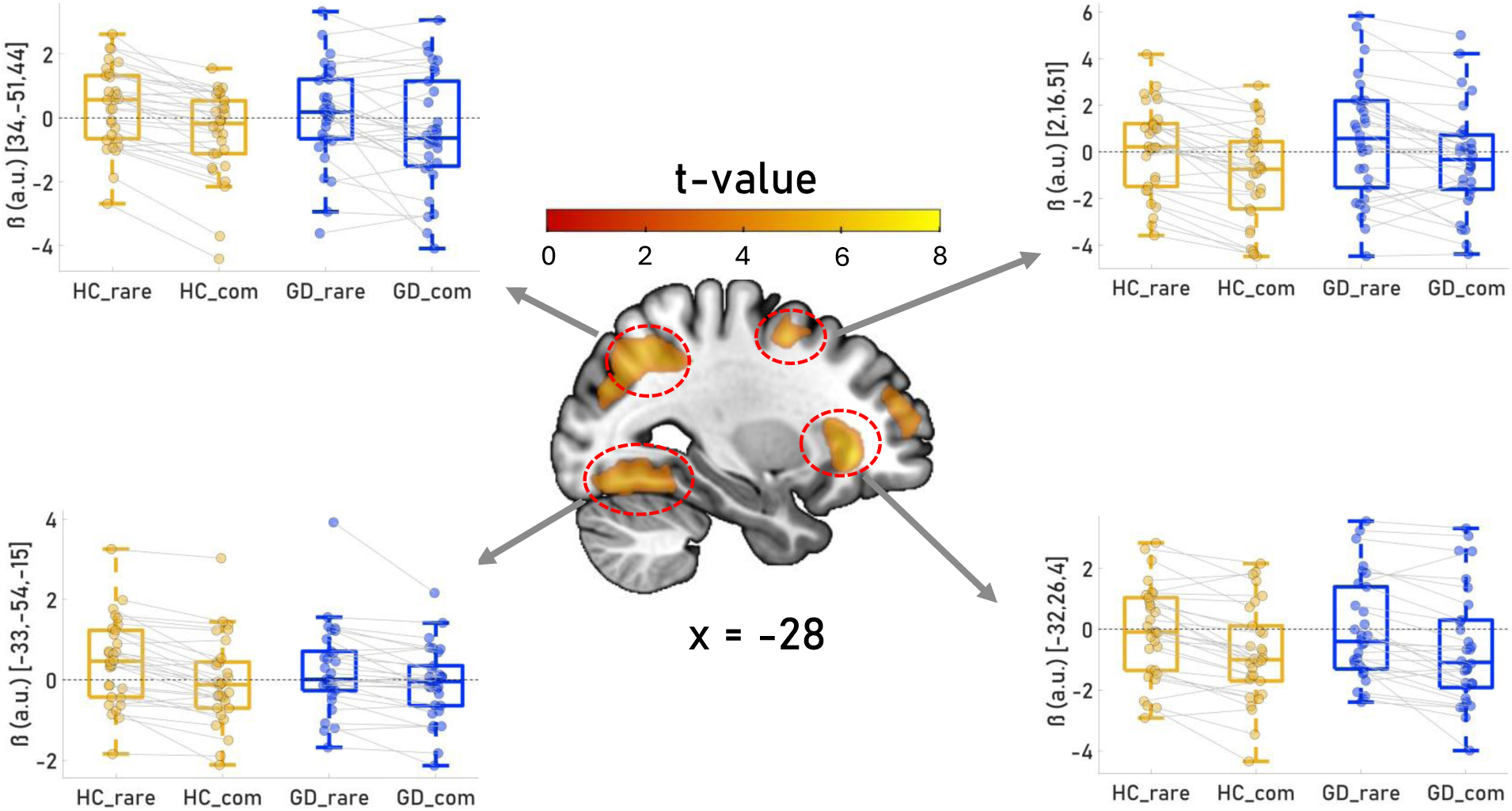
Neuronal correlates of task transition structure at second-stage onset. Whole-brain activation map showing the main effect of transition type (rare > common) as a parametric modulator at the onset of S2 (event regressor), across all participants (HC and GD). The map is thresholded at *p* < 0.001 (uncorrected) and reveals significant activation in multiple brain regions sensitive to the transition structure of the task. left inferior frontal gyrus [x = -32, y = 26, z = 4], dorsomedial prefrontal cortex [x = 2, y = 16, z = 51], right inferior parietal lobule [x = 34, y = -51, z = 44]; cerebellum [-33, -54, -15]; Insets in the upper corners display beta estimates extracted from four peak voxels within relevant activation clusters. Individual data points are overlaid on group-wise boxplots, with data from HC shown in yellow and GD in blue. a.u. = arbitrary units; No significant group differences were observed at these peaks.

#### Prediction Error and Reward Effects (GLM1, GLM2)

Neural correlates of model-free (MF) and model-based (MB) prediction errors (PEs) at S2 onset, were examined using a flexible factorial model with the factors group (HC vs. GD) and PE type (MF vs. MB), focusing on a reward region of interest (ROI) (Rangel lab: http://www.rnl.caltech.edu/resources/index.html), including ventral striatum, vmPFC, PCC and ACC. However, within this ROI, no voxels showed significant associations with either MF or MB prediction errors (SVC, FWE-corrected at p < 0.05). Similarly, a whole-brain analysis did not reveal significant main effects for either prediction error type across groups.

GD individuals exhibited significantly stronger model-based PE signals than HC in the medial frontal gyrus: x/y/z = 2/10/0), z-value = 5.23, SVC-corrected) and in the left superior frontal gyrus: x/y/z = -6/36/4, z = 5.00, SVC-corrected). No other significant group differences were observed for either MF or MB prediction error signals.

We next examined neural responses to reward at the outcome stage (GLM1/GLM2). In GLM1 we included a regressor for outcome onset combined with a parametric modulator reflecting the MF PE. We additionally used GLM2, in which outcome trials were split into two separate regressors based on relative reward magnitude (high vs. low, see methods). Both analyses focused the above-mentioned reward ROI mask. For Reward PE, we observed no significant effects in this ROI, analysis of high versus low reward trials revealed reward -related effects in the left posterior cingulate cortex (peak coordinates (x = -3, y = -34, z = 34), z-value = 5.75, p < 0.001), the left superior frontal gyrus (SFG; peak coordinates (x = -8, y = 58, z = -2), z-value = 5.50, p < 0.001), left ventral striatum (peak coordinates (x = 12, y = 10, z = -12), z-value = 4.05, p = 0.019) and right ventral striatum (peak coordinates (-12, 10, -10), z-value = 3.73, p = 0.056), (Figure 10; Supplementary Table S7). Again, no group differences were observed with relevant ROIs and across the whole brain.

**Figure 10:**
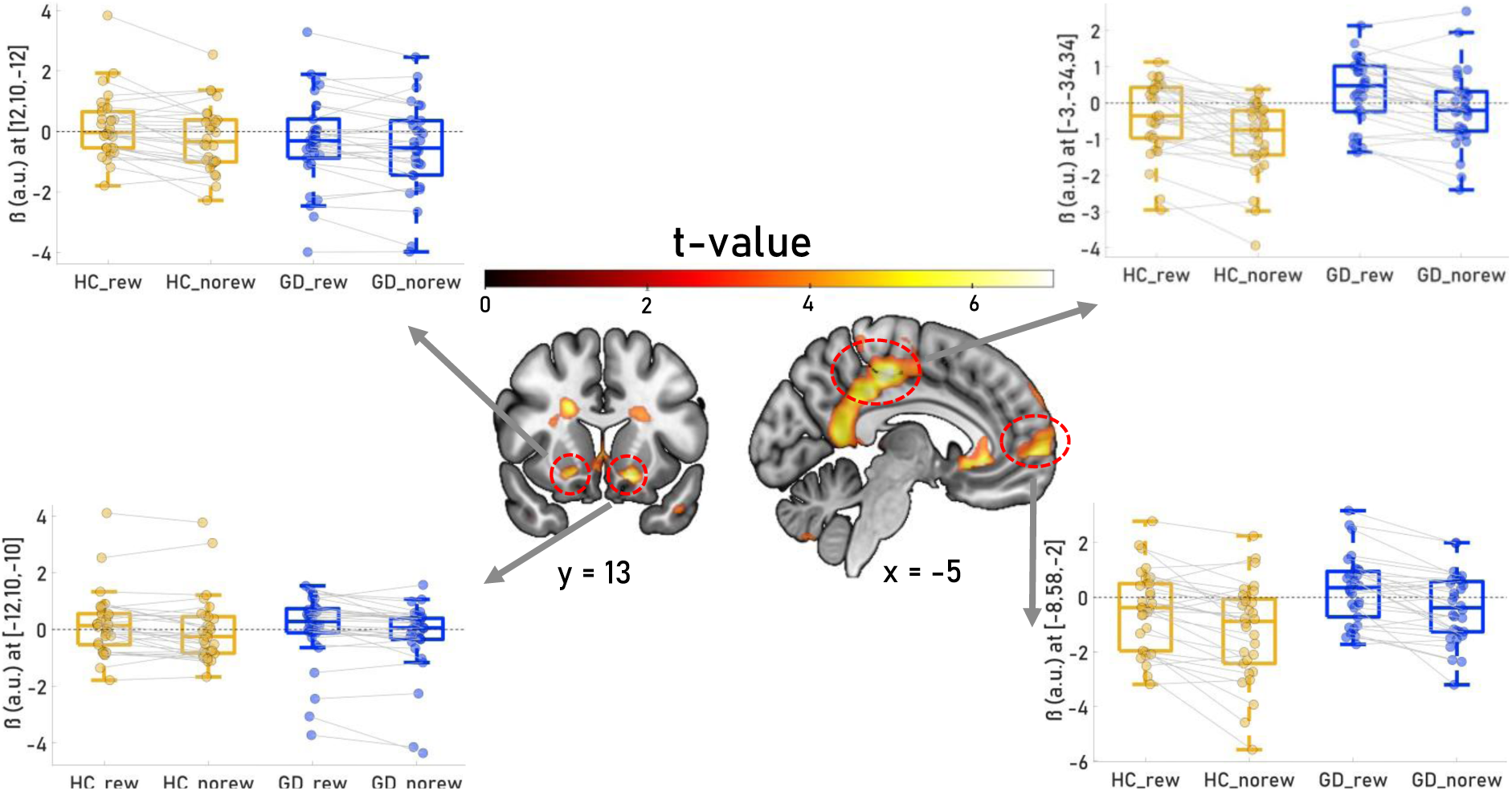
Brain regions showing significant activation for high vs. low reward contrasts. The central panel displays a contrast image illustrating regions with increased BOLD activation in response to high vs. low rewards. Significant activations were observed in the left superior frontal gyrus (SFG) (MNI: [- 8, 58, -2]), right ventral striatum (putamen and caudate, MNI: [12, 10, -12]), right superior frontal gyrus (SFG) (MNI: [9, 16, -4]), and the left ventral striatum (putamen and caudate, MNI: [-12, 10, -10],); The map is thresholded at p < 0.001 (uncorrected). In the upper and lower corners, beta weights from both HC (yellow) and GD individuals (blue) are shown as dots overlaid on boxplots for each region. a.u. = arbitrary units.

#### Exploratory Analysis of S1 Effects (GLM3)

Since neither group differences in S2 transition effects (‘state prediction errors’) nor MB prediction errors at S2 could account for the reduced MB control in the GD group, we next explored potential differences in S1 effects in more detail. GML3 included distinct S1 regressors based on previous reward (high vs. low), and previous transition (common vs. rare), resulting in four event types: (1) high reward + common transition (rc), (2) high reward + rare transition (rr), (3) low reward + common transition (uc), and (4) low reward + rare transition (ur). We reasoned that group differences in the transition structure model might already manifest at S1 onset. For example, if a participant received a low reward following a common transition, or a high reward following a rare transition, the next S1 onset might involve a strategy adjustment via executive control circuits. Following this rationale, we constructed the following contrasts for each participant in the first-level model: (S1 onset_rr_ + S1 onset_uc_) > (S1 onset_rc_ + S1 onset_ur_). These contrasts were compared between groups using a two-sample t-test. We indeed observed robust effects linked to behavioral adjustment in right (DLPFC) (MNI: [45, 34, 28], z = 5.06) and the right inferior parietal lobule (IPL) (MNI: [48, -51, 42], z = 4.35) (see Figure 11). However, again, no group differences were observed.

**Figure 11:**
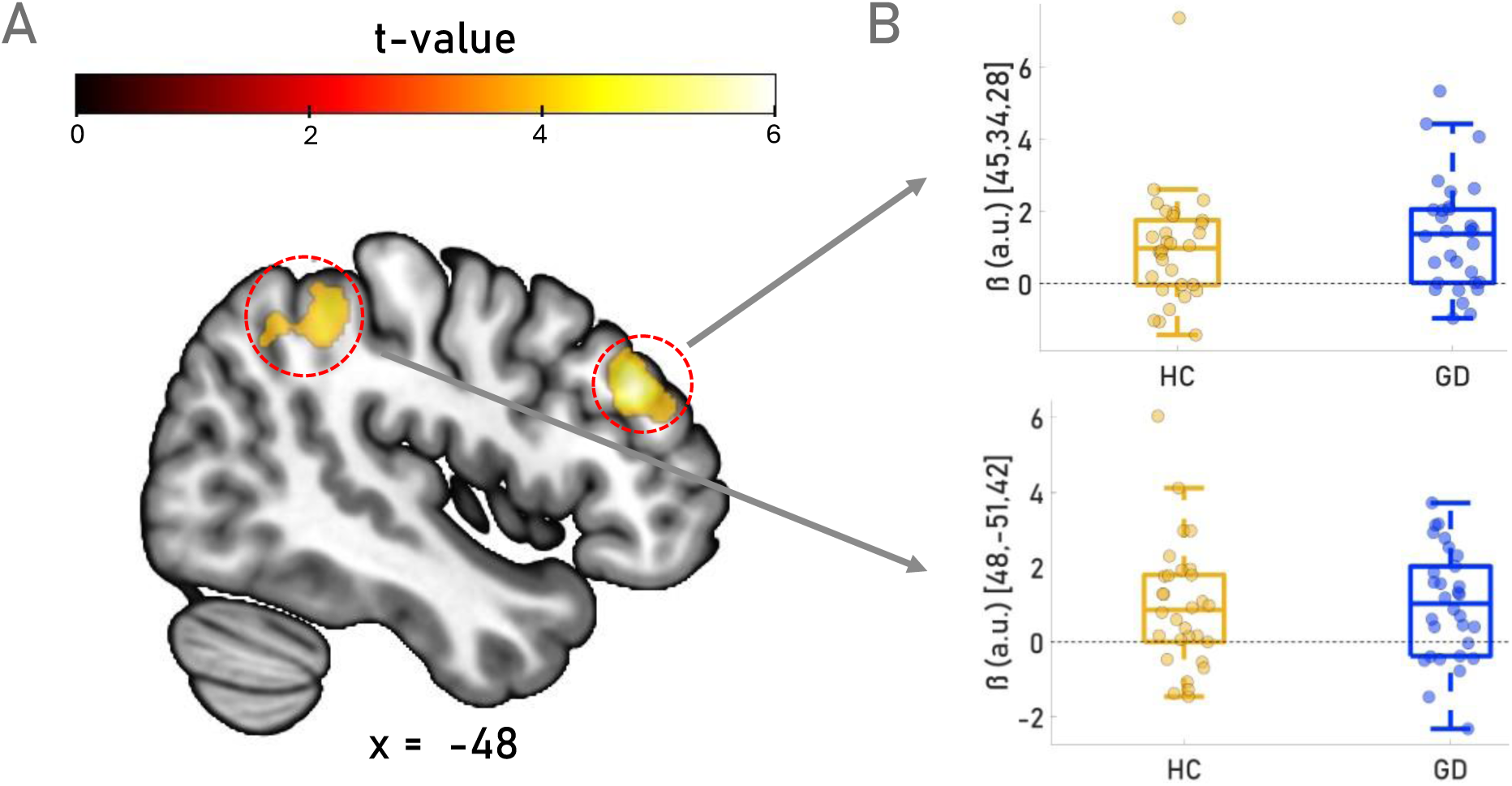
Differential BOLD responses at Stage 1 onset for events that might foster potential behavioral adjustments (i.e. choice switches). **A:** Results from the exploratory GLM contrasting the events (S1 onset_rr_ + S1 onset_uc_) > (S1 onset_rc_ + S1 onset_ur_) across both experimental groups (HC and GD). Significant activations were observed in the right dorsolateral prefrontal cortex (DLPFC) (MNI: [45, 34, 28], *z* = 5.06) and the right inferior parietal lobule (IPL) (MNI: [48, -51, 42], *z* = 4.35) during events that were hypothesized to require a behavioral adjustment compared to events where such an adjustment was not expected. **B:** Beta estimates extracted from the peak voxel in the PCC and DLPFC. Boxplots depict the distribution of parameter estimates for GD (blue) and HC (yellow). A.U. = arbitrary units; No group differences were found between GD and HC. The map is thresholded at p < 0.001 (uncorrected).

#### Neural Prospection Vividness Signature Predicts Model-Based Control

In a last exploratory analysis, we linked model-based control (ß_mb_-parameter) to potential fMRI signatures of *vivid prospection.* This analysis was motivated by the idea that model-based control depends on internal simulations of future outcomes, a cognitive process that might be linked to with vivid prospection. We reasoned greater reliance on MB control might be associated with a more vivid imagery of future outcomes. To this end, we applied a recently developed whole-brain decoder for prospection vividness (Lee et al., 2022) to the S1 onset contrast images to derive a vividness scores for each participant (see methods section for details).

Across all participants, there was a significant correlation of MB control with the neural vividness measure (*r* = 0.3, p = 0.018, see Figure 12). This correlation was primarily driven by the HC group (*r* = 0.46, p = 0.0097; see Figure 12), whereas the relationship was numerically attenuated in the GD group (*r* = 0.25, p = 0.18) (although correlations did not differ significantly between groups, z= 0.91, p=.36). This suggests that vivid imagery may generally support model-based control, irrespective of group membership.

**Figure 12.**
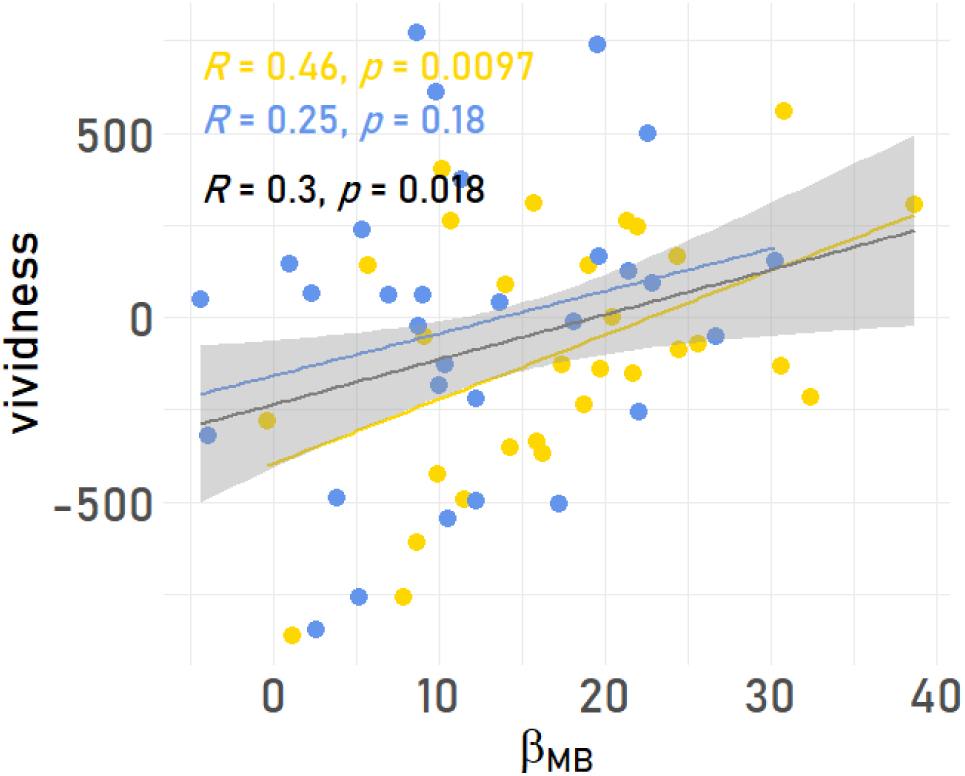
Association Of Model-Based Control and Neural Marker for Vivid Prospection. Yellow dots represent healthy controls, and blue dots represent individuals with gambling disorder. Regression lines and correlation coefficients (R) are shown for each group and across all participants. A significant positive correlation was observed in healthy controls (R = 0.46, p = 0.0097), while the correlation in GD individuals was weaker and not statistically significant (R = 0.25, p = 0.18). When data from both groups were pooled, a modest but significant correlation was found (R = 0.3, p = 0.018). Shaded regions represent 95% confidence intervals.

## Discussion

We examined the neurocomputational mechanisms underlying impaired model-based control in individuals exhibiting symptoms of disordered gambling (GD) and matched health controls (HC). Using a modified sequential reinforcement learning task (two-step-task), we replicated previous findings of reduced performance in the GD group. Computational modelling using standard softmax action selection replicated a substantial reduction in MB control in the GD group. Reinforcement learning drift diffusion models (DDMs) revealed that this MB control deficit in the GD group was linked to changes across several computational mechanisms (see discussion below). Despite these robust behavioural and computational differences, fMRI results were largely similar between groups across both task stages. For example, both groups showed robust fronto-parietal activation in S2 following rare vs. commong transitions (similar to state prediction errors, see Gläscher et al., 2010), and reward-related effects in striatal and ventromedial prefrontal areas were similar between groups. Likewise, both groups exhibited increased hippocampal activation in S1 following high reward feedback, and increased dmPFC activation in S1 following rare transitions, indicating sensitivity to both prior outcomes and task structure. An exploratory analysis revealed a positive relationship between MB control and neural signatures of vivid prospection, suggesting that mental imagery may support MB control.

Model-agnostic and model-based results replicated previous findings of reduced MB control in problem gambling (Wyckmans et al., 2019, Bruder et al., 2021). Deficits in goal-directed control of have been consistently hypothesised to represent a crucial component of habitual and compulsive behavior in addiction (Everitt & Robbins, 2005; Voon et al., 2015; 2017). GD has likewise been linked to compulsivity phenotypes (van Timmeren et al. 2018). Such inflexibility may foster maladaptive behaviours despite negative consequences in addiction (Voon et al., 2015; Gillan et al., 2016). In line with these considerations, our computational modelling approach extended standard hybrid TST models by taking into account higher-order perseveration (HOP) behaviour (Miller et al., 2019; Collins & Cockburn, 2020; Palminteri, 2023; Brands et al., 2025). The inclusion of a “value-free” perseveration term improved model fit across both groups, and this mechanism was also incorporated into the DDMs. The additional HOP step-size parameter from the softmax model differed between groups, such that GD participants tended to show perseveration over longer time scales. This pattern is in line with a diminished capacity for behavioural adaptation in changing environments (i.e. transition and reward structure of the task; Fineberg et al., 2014; van Timmeren et al., 2018; Perandrés-Gómez et al., 2021). In the present task, a higher updating rate (i.e. shorter tail, more FOP-like behaviour) as seen in HC participants, may have enabled more flexible behaviour.

We then used combined reinforcement learning DDM variants of the hybrid RL model to leverage both choices and RTs in model estimation. This confirmed a superior for of the DDM_SIGMOID_ in both groups (Fontanesi et al., 2019; Peters & D’Esposito, 2020), and model performance was confirmed via posterior predictive checks. Interestingly, compared to the softmax models, in the DDM_SIGMOID_, performance deficits in the GD group were linked to a more complex pattern of parameter changes. Here, 𝑣_*MB*_ (the model-based weight on the drift rate) showed only a marginal reduction in the GD group. In contrast, the GD group exhibited reduced maximum drift rates (*ν*_*MAX*_) and non-decision times (*τ*) in both task stages. Both *ν*_*MAX* −*S*1_ and *τ*_*S*1_were positively correlated with *β*_*MB*_ from the softmax model, highlighting that the MB deficit in GD was accounted for by changes across several parameters in the DDM_SIGMOID_. The finding of lower non-decision times in both stages in the GD group dovetails with previous reports (Wiehler & Peters, 2024) and may also reflect higher motor impulsivity and/or urgency in GD (Billieux et al., 2012; Kräplin et al., 2014; Chowdhury et al., 2017). Overall RTs were not significantly different between groups, showing that DDM-based decomposition can reveal subtle group differences that are overlooked in standard RT analyses (see also Wiehler & Peters, 2024). Posterior predictive checks of the DDM_SIGMOID_ further revealed a more nuanced insight into performance differences between groups, especially in S2. Here, the GD group exhibited a higher proportion of optimal choices at low value contrasts compared to the HC group (i.e. high difficulty; Figure 2C, 2D). However, GD participants did not substantially benefit from further increases in value contrast, while HC participants did, consistent with the overall superior performance in this group. This pattern is again in line with an important role of behavioural inflexibility (i.e. lack of dynamic adaptation) in GD (Rochat et al., 2019; Winstanley & Clark, 2016; Voon et al., 2017).

Correlation analyses revealed no associations between the HOP step-size and MB control parameters with clinical gambling-related scales in the GD group. However, gambling-related cognitions (GRCS, Raylu & Oei, 2004) were negatively associated with the HOP step-size parameter α_HOP_, such that lower scores, were associated with higher α_HOP_ estimates. That is, α_HOP_ was more similar to parameter estimates in the control group with decreasing GRCS total scores, suggesting a potential link between maladaptive perseveration behaviour and erroneous gambling-related cognitions. However, further investigations using more detailed and extensive assessment of clinical features and/or cognitive distortions are warranted in larger and more diverse populations.

Despite the robust group differences in performance and model parameters, neuroimaging findings were characterized by overall very similar patterns of effects in the two groups. In both groups, activity at S1 onset in cortical and subcortical circuits, including bilateral hippocampus, was modulated by previous trial reward magnitude. This aligns with recent work (Bakkour et al., 2019) demonstrating that hippocampal activation tracks deliberation time in value-based choices, and may implicate memory retrieval and/or simulation in early evaluative processing. The idea that participants may have drawn upon memory- or prospection-based representations to guide upcoming decisions is also supported by our exploratory analysis of neural prospection vividness signatures in S1. This this end, we applied a previously published whole brain decoder (Lee et al. 2022) to S1 onset contrast images to derive neural vividness scores for each participant. Across all participants, this revealed a positive correlation of vividness scores with *β*_*MB*_, supporting the general idea that prospection may contribute to model-based control (Doll et a. 2015). Although this correlation was more pronounced in the control group, correlations did not differ significantly between groups, such that we are reluctant to draw additional conclusions at this point. A direct comparison of neural vividness indices also did not reveal significant group differences, arguing against the idea that group differences in prospection underlie the MB control deficits in the GD group.

We next examined neural sensitivity to task structure both with respect to S1 and S2 effects. For S1, BOLD responses were modelled as a function of the previous trial’s transition type (rare > common). Across both groups, robust activation was observed in the amPFC, with anterior PFC regions having previously been implicated in model-based control (Doll et al., 2015). Again, no significant group differences were observed. For S1, onset-related effects focused on transition type (rare vs. common), akin to analyses of state prediction errors (SPEs; Gläscher et al., 2010). In the context of multi-step RL, SPEs reflect the degree to which observed state transitions deviate from expectation (e.g. experiencing a rare transition when a common transition is anticipated). SPEs are central to model-based learning, as they allow an update of the agent’s internal model of the environment and may guide future decision-making. In both groups, rare transitions elicited robust activation in a distributed fronto-parietal control network closely aligned with previously described SPE-related networks (e.g., Gläscher et al., 2010). There was no credible evidence for group differences in these effects, arguing against the idea that impaired representations of the basic task transition structure underlie the MB control deficits in the GD group.

Contrary to our expectations and to previous work (e.g. Daw et al., 2011; Deserno et al., 2015; Kroemer et al., 2019), neither model-free nor model-based PEs at second-stage onset were significantly associated with BOLD activity within the canonical reward network (including ventral striatum and vmPFC) when analysed across all participants or within groups. One potential reason for this lack of modulation may be that, in contrast to the work cited above, the modified TST version employed here used continuous rewards, rather than binary rewards. This may have attenuated neural prediction error related effects, which may be more salient for categorical vs. continuous feedback. This change in reward feedback may have resulted in participants relying less on prediction error-driven updating and more on prospective, simulation-based mechanisms. The results from our exploratory analysis of neural vividness signatures partly supports this idea. However, the lack of PE-related effects may also be due to the clinical and matched control samples tested here, which contrasts with WEIRD samples from some previous studies. Based on this lack of PE-related neural effects, we also assessed categorical reward effects, contrasting high vs. low reward feedback (see methods). This revealed robust effects in canonical reward-related regions, including bilateral ventral striatum and medial prefrontal regions, highlighting that the lack of PE-related effects was not due to e.g. signal loss, and is unlikely to be due to general lack of task engagement. Again, no significant group differences emerged in these effects, arguing against the idea that reduced reward related activity underlies the MB control deficit in the GD group (Clark et al., 2019).

Finally, an exploratory analysis assessed whether BOLD responses at S1 onset varied with the preceding trial’s outcome and transition type. This contrast aimed to capture neural markers of model-based behavioural adjustment (see Kroemer et al., 2019), mirroring the behavioural analysis of stay probabilities. We observed greater activation in right dorsolateral prefrontal cortex (DLPFC) and right inferior parietal lobule occurred following rewarded rare and unrewarded common trials, i.e. in conditions associated with behavioural adjustment in model-based frameworks. These regions are linked to cognitive control and strategy adjustment (Badre & D’Esposito, 2009; O’Reilly, 2010), but given the exploratory nature of this analysis, we are hesitant to draw firm conclusions regarding their precise functional role. Nonetheless, there were no credible group differences in these effects, which may reflect preserved neural signatures of model-based behavioural adjustment in S1. In line with the robust transition-related effects on activity related to S2 onset in both groups, this suggests that impairments in GD may reflect difficulties in consistently applying knowledge about the task structure, rather than a failure to understand it.

Taken together, although the GD group exhibited a clear deficit in model-based control (resulting in reduced performance and corresponding changes in computational model parameters), fMRI analyses revealed no credible corresponding group differences. We replicated core effects related to reward outcome and task transition structure representations, confirming that our design was overall sensitive to reveal these effects. What to make of this discrepancy? The null findings regarding group differences may reflect high interindividual variability in how participants implement and represent model-based strategies at the neural level, which may have diluted group-level effects, especially in the absence of large sample sizes or more targeted, individually tailored analytical approaches. Intact neural sensitivity to task structure in core networks implicated in SPEs across both groups suggests that GD individuals process task structure in a manner similar to HC. Behavioural impairments may therefore stem from a deficit in consistently applying this knowledge to guide choices in a flexible and adaptive manner (however, neural indices of behavioural adjustment during S1 choices in PFC areas were also not credibly different between groups). It might also be speculated that more subtle group differences lie in network-level interactions that were not captured by the present analysis. Future studies using time-resolved or connectivity-based methods may provide deeper insight into the neural mechanisms underlying MB control impairments in GD.

Several limitations of the present study need to be addressed. First, our sample consisted of only male participants (reflecting prevalence differences). Future studies would benefit from a consideration of potential gender effects. Second, as mentioned above, we did not examine functional connectivity differences between groups. GD has been linked to changes in functional coupling in limbic circuits (Peters et al., 2013), which may be a fruitful target for future investigations. Third, while our sample size is in line with previous work and was based on an *a priori* power analysis, it is still too small to comprehensively address individual differences in gambling subtypes (Nower et al., 2021) and preferred gambling formats (our GD sample consisted of individuals engaging in various types of online and terrestrial gambling). Finally, while we carried out comprehensive model comparison and posterior predictive checks for model validation, conclusions drawn from computational modelling are inherently restricted to the specific models included in the model space. E.g., participants may have employed alternative “model-based” strategies not captured by the models employed here (Da Silva & Hare, 2020). Nonetheless, the behavioural impairment in the GD group was also evident across several model-agnostic measures (stay probability, overall performance, S2 RT effect).

Taken together, our results substantiate previous findings of MB control impairments in GD (see e.g. Bruder et al., 2021; Wyckmans et al., 2019), and demonstrate that this impairment is likely linked maladaptive adjustments across multiple computational mechanisms (higher-order perseveration, reduced MB control), illustrating the complexity of MC control impairments in GD. The lack of credible neural group differences, despite a replication of core task-related effects in our design, highlights the need for more nuanced neurocognitive models that integrate multiple decision-making systems.

## Supplementary Information

**Table S1.** Linear mixed model results for S1 response times (RTs)

|  | estimate | SE | t-Value | p-Value |
| --- | --- | --- | --- | --- |
| **intercept** | **0.65** | **0.01** | **47.82** | **<.001** |
| reward | -0.003 | 0.002 | -1.23 | .224 |
| transition | -0.001 | 0.01 | -0.52 | .604 |
| group | -0.007 | 0.002 | -0.54 | .590 |
| reward*transition | 0.001 | 0.002 | 0.60 | .550 |
| **reward\*group** | **-0.005** | **0.002** | **-2.24** | **.029** |
| transition*group | 0.003 | 0.002 | 1.21 | .231 |
| reward*transition*group | 0.004 | 0.002 | 1.51 | .138 |

**Table S2.** Linear mixed model results for S1 stay probabilities p(stay)

|  | estimate | SE | z | p-Value |
| --- | --- | --- | --- | --- |
| **Intercept** | **1.14** | **0.11** | **10.52** | **<.001** |
| **reward** | **0.10** | **0.03** | **3.36** | **<.001** |
| transition | -0.04 | 0.03 | -1.67 | .095 |
| **group** | **-0.42** | **0.12** | **-3.48** | **<.001** |
| BDI | 0.17 | 0.09 | 1.91 | .057 |
| BIS | -0.07 | 0.08 | -0.81 | .419 |
| FTND | 0.16 | 0.09 | 1.71 | .088 |
| AUDIT | -0.08 | 0.08 | 0.92 | .360 |
| **reward\*transition** | **0.38** | **0.05** | **8.17** | **<.001** |
| reward*group | 0.03 | 0.03 | 0.99 | .321 |
| transition*group | 0.01 | 0.02 | 0.42 | .671 |
| **reward\*transition\*group** | **-0.10** | **0.05** | **-2.22** | **.027** |

**Table S3.** Linear mixed model results for S2 response times (RTs)

|  | estimate | SE | t-Value | p-Value |
| --- | --- | --- | --- | --- |
| **intercept** | **0.75** | **0.01** | **54.09** | **<.001** |
| **transition** | **-0.001** | **0.01** | **-8.85** | **<.001** |
| group | -0.007 | 0.002 | -0.23 | .817 |
| **transition\*group** | **0.003** | **0.002** | **2.48** | **.016** |

**Table S4.**
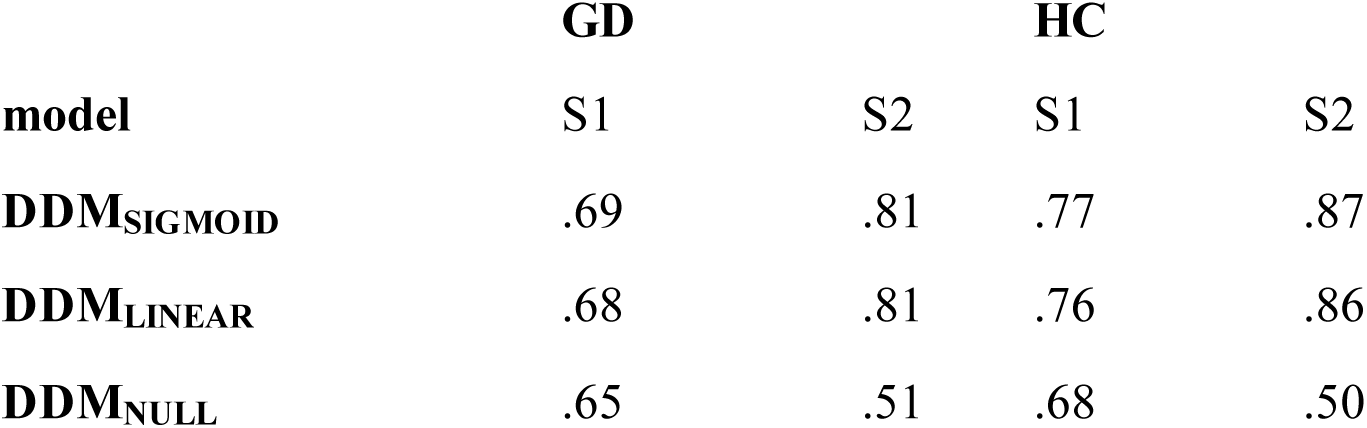
Mean Proportion of Correct Choice Predictions per DDM, Group, & Task-Stage.

|  | **GD** |  | **HC** |  |
| --- | --- | --- | --- | --- |
| **model** | S1 | S2 | S1 | S2 |
| **DDM<sub>SIGMOID</sub>** | .69 | .81 | .77 | .87 |
| **DDM<sub>LINEAR</sub>** | .68 | .81 | .76 | .86 |
| **DDM<sub>NULL</sub>** | .65 | .51 | .68 | .50 |

**Table S5.** Association of model parameters reflecting MB control and clinical measures.

| DV | Predictor | Estimate | SE | t-value | p |
| --- | --- | --- | --- | --- | --- |
| **$\beta_{MB}$** | Intercept | 15.58 | 9.20 | 1.69 | .10 |
|  | GD score | -0.21 | 2.90 | -0.07 | .94 |
|  | GRCS | -0.35 | 0.33 | -1.08 | .29 |
|  | BDI | -0.04 | 0.21 | -0.19 | .85 |
|  | FTND | 0.26 | 0.67 | 0.39 | .70 |
|  | BIS | -0.14 | 0.26 | -0.52 | .60 |
|  | AUDIT | 0.14 | 0.23 | 0.63 | .54 |
| $\nu_{MB}$ | Intercept | 15.86 | 12.20 | 1.30 | .21 |
|  | GD score | 0.37 | 3.85 | 0.10 | .92 |
|  | GRCS | -0.40 | 0.43 | -0.92 | .36 |
|  | BDI | -0.13 | 0.28 | -0.45 | .66 |
|  | FTND | 0.95 | 0.89 | 1.06 | .30 |
|  | BIS | -0.00 | 0.34 | -0.01 | .99 |
|  | AUDIT | 0.22 | 0.30 | 0.72 | .48 |

Note. GD-score: gambling severity score. GRCS: gambling related cognitions; BDI: depression; FTND: nicotine dependence; BIS: impulsivity; AUDIT: alcohol dependence. $\beta_{MB}$ : model-based control parameter from the softmax model. $\nu_{MB}$ : model-based effect on the drift rate from the DDM<sub>SIGMOID</sub> .

**Table S6.** Regression analysis of model parameters and self-report scales.

| DV | Predictor | Estimate | SE | t-value | p |
| --- | --- | --- | --- | --- | --- |
| $\alpha_{HOP}$ | Intercept | 0.39 | 0.44 | 0.89 | .38 |
|  | GD-score | -0.10 | 0.14 | -0.69 | .49 |
|  | **GRCS** | **-0.03** | **0.02** | **-2.33** | **.03** |
|  | BDI | 0.00 | 0.01 | 0.40 | .70 |
|  | FTND | -0.04 | 0.003 | -1.26 | .22 |
|  | BIS | 0.02 | 0.01 | 1.54 | .14 |
|  | AUDIT | -0.01 | 0.01 | -1.04 | .31 |
| $\alpha_{HOP-DDM}$ | Intercept | 0.48 | 0.35 | 1.38 | .18 |
|  | GD-score | -0.11 | 0.11 | -1.04 | .31 |
|  | **GRCS** | **-0.03** | **0.01** | **-2.58** | **.02** |
|  | BDI | -0.00 | 0.01 | -0.05 | .95 |
|  | FTND | -0.03 | 0.03 | -1.21 | .24 |
|  | **BIS** | **0.02** | **0.01** | **2.07** | **.05** |
|  | AUDIT | -0.01 | 0.01 | -0.80 | .43 |

Note. GD-score: gambling severity score. GRCS: gambling related cognitions; BDI: depression; FTND: nicotine dependence; BIS: impulsivity; AUDIT: alcohol dependence. $\alpha_{HOP}$ : HOP step-size parameter from the model using softmax action selection. $\alpha_{HOP-DDM}$ : DDM HOP step-size parameter.

**Table S7:**
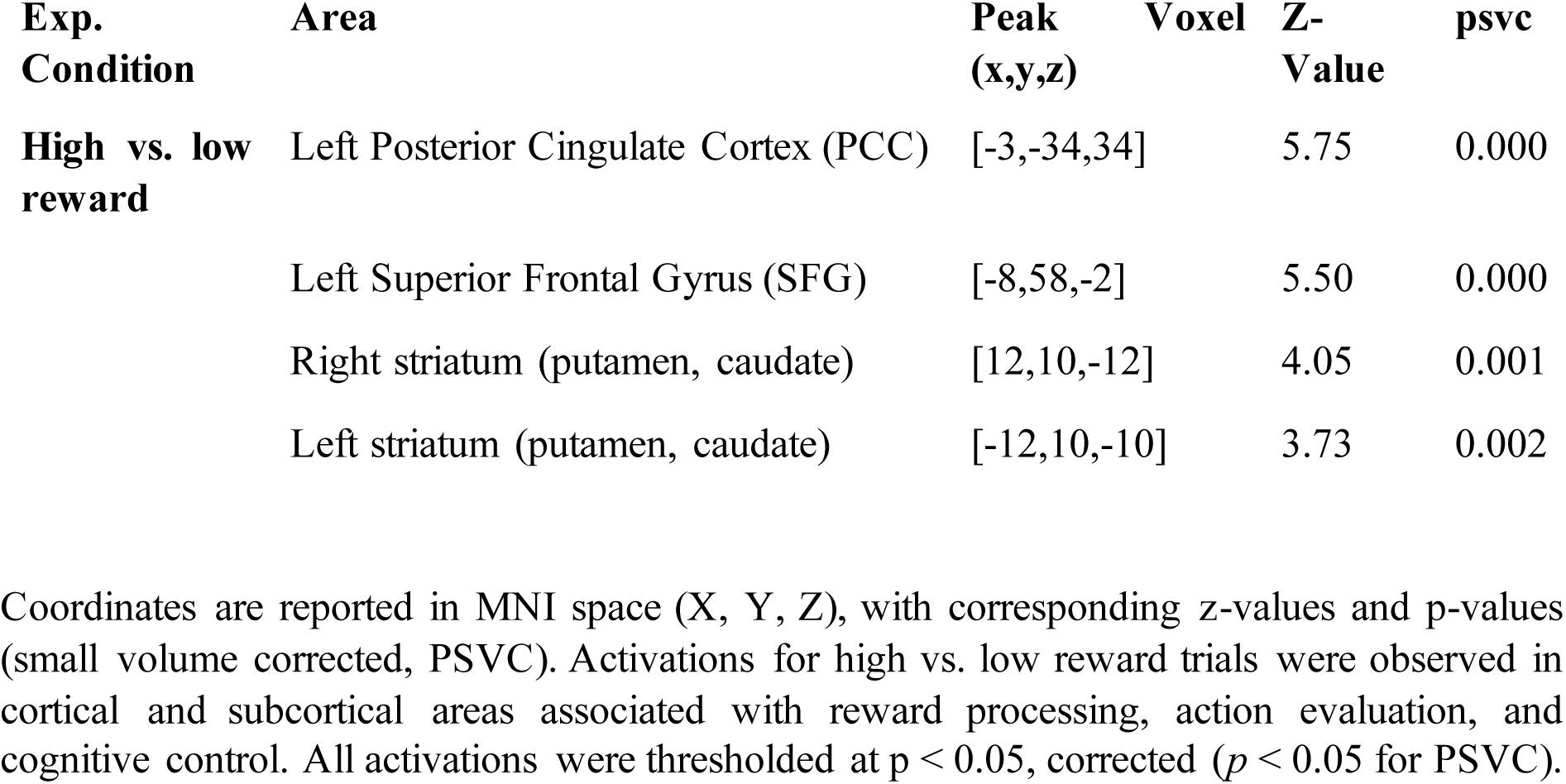
Brain regions showing significant activation for high vs. low reward contrasts during the outcome stage.

